# Neural crest cell derived DKK1 modulates Wnt signalling in the second heart field to orchestrate cardiac outflow tract development

**DOI:** 10.1101/2025.03.27.645622

**Authors:** Sophie Wiszniak, Dimuthu Alankarage, Iman Lohraseb, Ceilidh Marchant, Genevieve Secker, Wendy Parker, John Toubia, Melissa White, Sandra Piltz, Markus Tondl, Eleni Giannoulatou, David Winlaw, Gillian M. Blue, Congenital Heart Disease Synergy Group, Patrick P. L. Tam, Paul Thomas, Natasha Harvey, Sally L. Dunwoodie, Quenten Schwarz

**Affiliations:** Centre for Cancer Biology, University of South Australia and SA Pathology, Adelaide, SA, 5000, Australia; Victor Chang Cardiac Research Institute, Darlinghurst, NSW, 2010, Australia; University of New South Wales, Kensington, NSW, 2033, Australia; ACRF Cancer Genomics Facility, SA Pathology, Adelaide, SA, 5000, Australia; University of Adelaide and South Australian Health and Medical Research Institute, Adelaide, SA, 5000, Australia; School of Clinical Medicine, Faculty of Medicine and Health, UNSW Sydney, NSW 2052, Australia; Heart Center, Ann and Robert H. Lurie Children’s Hospital of Chicago, Chicago, IL 60611, USA; Feinberg School of Medicine, Northwestern University, Chicago, IL 60611, USA; Heart Centre for Children, The Children’s Hospital at Westmead, Westmead, NSW, 2145, Australia; Sydney Medical School, Faculty of Medicine and Health, University of Sydney, Sydney, NSW, 2000, Australia; Children’s Medical Research Institute, University of Sydney, Westmead, NSW, 2145, Australia; School of Medical Sciences, Faculty of Medicine and Health, UNSW Sydney, NSW 2000, Australia

## Abstract

Heart morphogenesis is highly complex, and depends on the generation of diverse cell types which interact with each other in an orchestrated manner to remodel the primitive heart tube into a functional organ. Cardiac outflow tract formation critically depends on continued contribution of cardiac progenitor cells from the anterior second heart field to ensure proper growth of the outflow tract. Prior to entering the outflow tract, neural crest cells migrate in close apposition to the second heart field and may play important roles in regulating second heart field growth dynamics, however the molecular mechanisms by which neural crest cells interact with the second heart field have remained elusive. Here, we discover that neural crest cells are a primary source of Dickkopf1 (DKK1), a secreted Wnt signalling inhibitor, which modulates Wnt signalling activity in the second heart field to impose a balance between progenitor maintenance and differentiation. Further, we identify the ubiquitin ligase NEDD4 as a critical regulator of DKK1 levels, with disruption of *Nedd4* activity leading to outflow tract defects. In the context of disease pathogenicity, we show a novel human congenital heart disease variant of *NEDD4* has lost the ability to ubiquitinate DKK1, and is associated with heart defects in a mouse model of the genetic variant. Our findings point to an unexpected role for neural crest cells acting as a rheostat of Wnt signalling activity in cardiac progenitors, identifying a new molecular pathway underpinning correct outflow tract morphogenesis, and a new causative factor of congenital heart disease.

## Introduction

The cardiac outflow tract of the heart provides the pathway for blood to leave the ventricles and circulate to the body via the aorta, or to the lungs via the pulmonary artery. Outflow tract development is complex and requires the contribution of cell types from multiple embryological origins, including second heart field cardiac progenitor cells, cardiomyocytes, smooth muscle, endocardium and neural crest cells ^1^. During early development, the primitive heart tube expands through the progressive addition of cardiac progenitors from the anterior second heart field, which give rise to myocardium of the outflow tract and right ventricle. As the outflow tract lengthens, it repositions to overlie the future interventricular septum. Inadequate addition of second heart field cells to the outflow tract leads to insufficient lengthening, and underpins conotruncal defects ^2^.

In the heart, neural crest cells form structural components of the outflow tract valves, arteries, and conotruncal septum. Classical defects associated with disrupted cardiac neural crest cell development include persistent truncus arteriosus, where the outflow tract fails to septate into the aorta and pulmonary arteries ^3^. Prior to entering the outflow tract, neural crest cells migrate in close apposition to cardiac progenitors of the second heart field, and are suggested to play a role in modulating the growth dynamics of second heart field derivatives ^2^. However, prior models in which neural crest cells have been surgically or genetically ablated^4–7^ have not been amenable to elucidate how neural crest cells interact with the second heart field to control its growth, differentiation and morphogenesis.

NEDD4 is an E3 ubiquitin ligase that targets specific protein substrates for ubiquitination to modulate protein function and/or turnover ^8^. Prior studies have determined a necessary role for *Nedd4* in heart development, with full knockout mouse models exhibiting DORV and endocardial cushion defects ^9^. However, the cell-type specific mechanisms by which *Nedd4* regulates cardiac development remained unexplored. Using conditional knockout mouse models, we determined an essential tissue-specific role for *Nedd4* in neural crest cells for correct outflow tract development. *Wnt1-Cre; Nedd4^fl/fl^*embryos exhibit outflow tract defects typically associated with failed addition of second heart field cells to the outflow tract, suggesting a novel role for *Nedd4* and cardiac neural crest cells in regulating second heart field development. Furthermore, we identify DKK1 as a previously unknown substrate of NEDD4-mediated ubiquitination, and find that this functionality is impaired in a human variant of NEDD4 associated with the outflow tract defect Tetralogy of Fallot. We propose NEDD4 controls DKK1 protein levels in neural crest cells, which in turn modulates Wnt signalling in the second heart field to balance cardiac progenitor maintenance versus myocardial differentiation, underpinning normal outflow tract morphogenesis, and defining a potential pathogenic mechanism underlying congenital heart disease.

## Results

### *Nedd4* is required in neural crest cells for cardiac outflow tract development

Prior studies have reported heart defects in *Nedd4^-/-^*embryos ^9^. We specifically assessed outflow tract defects in *Nedd4^-/-^* embryos, revealing defects at full penetrance with variable phenotypes, most commonly persistent truncus arteriosus (PTA) and transposition of the great arteries (TGA) (Fig. 1A,E,I-L,Q). To determine the cell types in which *Nedd4* activity is critically required, we conditionally removed *Nedd4* using tissue-specific Cre-drivers for cell lineages important in outflow tract development. Ablation of *Nedd4* in second heart field (*Mef2cAHF-Cre; Nedd4^fl/fl^*) and endothelial/endocardial cells (*Tie2-Cre; Nedd4^fl/fl^*) had no discernible impact on development of the outflow tract (Fig. 1C,D,G,H). In contrast, ablation of *Nedd4* in the neural crest cells (*Wnt1-Cre; Nedd4^fl/fl^*) led to fully penetrant outflow tract defects such as double-outlet right ventricle (DORV) and overriding aorta (Fig. 1B,F,M-P,Q). Importantly, the outflow tract defects identified in *Nedd4^-/-^* and *Wnt1-Cre; Nedd4^fl/fl^* embryos are considered artery-ventricle alignment type defects that present on a variable spectrum of severity. These findings point to a critical cell type-specific role for *Nedd4* in neural crest cells to orchestrate outflow tract development, consistent with previous demonstration of essential roles for *Nedd4* in neural crest cells in craniofacial morphogenesis and peripheral nervous system development ^10–12^.

**Figure 1.**
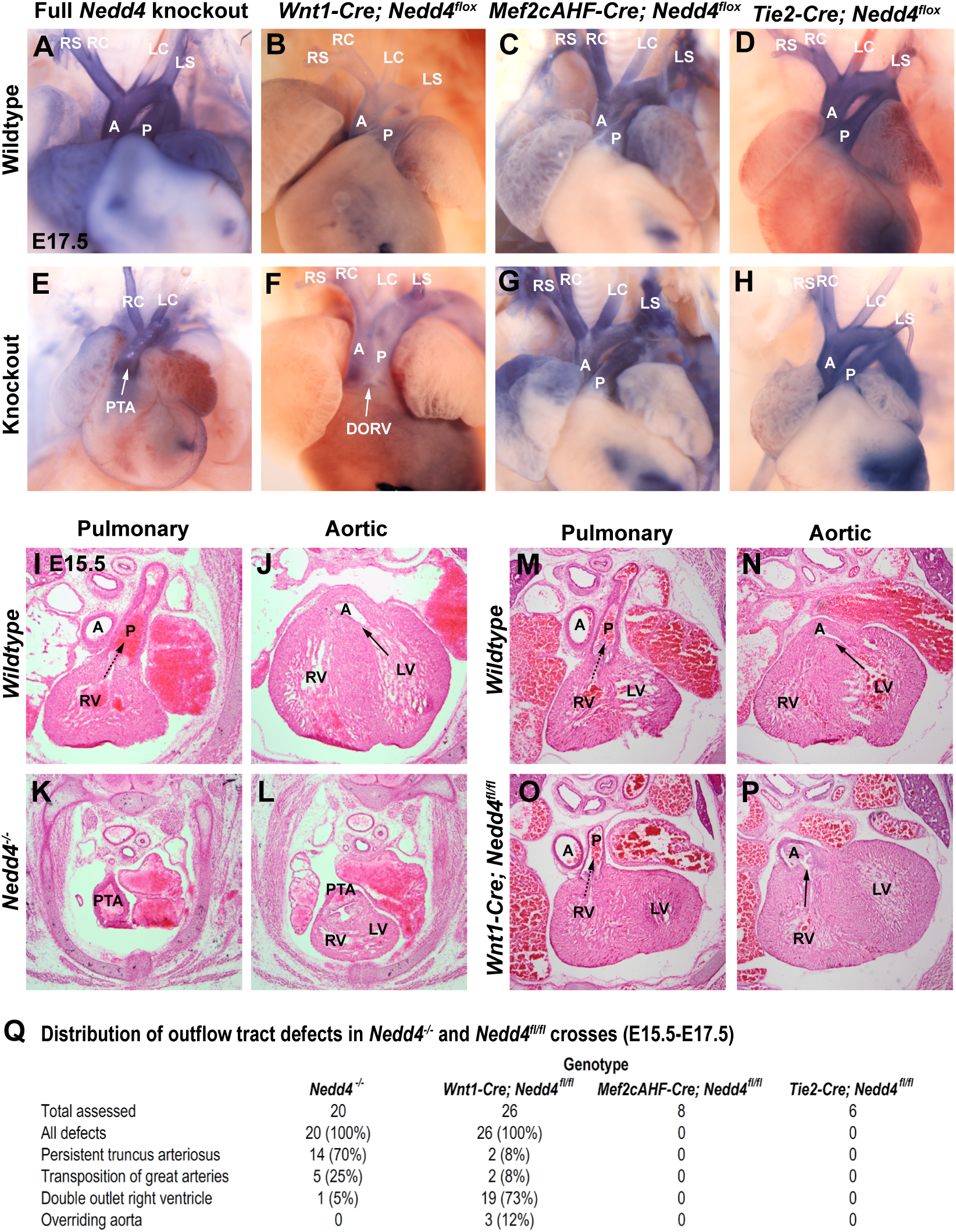
*Nedd4* activity in neural crest cells is required for cardiac outflow tract development. **A-H:** Whole E17.5 hearts from *Nedd4* full- or conditional-knockout mouse strains. Hearts were injected with Evans Blue solution in the right and left ventricles to aid visualisation of the outflow tract and branching arteries. A, aorta; P, pulmonary artery; RS, right subclavian artery; RC, right common carotid artery; LC, left common carotid artery; LS, left subclavian artery; PTA, persistent truncus arteriosus; DORV, double outlet right ventricle. **I-P:** Transverse sections through the heart and outflow tract of E15.5 embryos at the level of the pulmonary or aortic valve region from wildtype and *Nedd4^-/-^* strains (I-L), and from wildtype and *Wnt1-Cre; Nedd4^fl/fl^* strains (M-P), stained with H&E. Dashed arrow indicates ventricle patency with the pulmonary artery, and solid arrow indicates ventricle patency with the aorta. RV, right ventricle; LV, left ventricle. **Q:** Frequency of outflow tract defects in indicated mouse strains, classified by histological staining of tissue sections as in (I-P).

### Loss of *Nedd4* function in neural crest cells compromises outflow tract lengthening and elicits premature differentiation of the second heart field

The myocardial component of the outflow tract is derived from cardiac progenitors of the second heart field, with growth of the outflow tract dependent on the continued addition of cardiac progenitors to lengthen the outflow tract, allowing for heart looping and outflow tract rotation to establish correct artery-ventricle alignment ^1,2^. Underpinning the artery-ventricle alignment defects in *Wnt1-Cre; Nedd4^fl/fl^* embryos, defective outflow tract lengthening at E10.5 (Fig. 2A) led to incomplete clockwise rotation of the outflow tract at E12.5, and mispositioning of the aorta over the right ventricle (Fig. 2B). Such artery-ventricle alignment defects typically stem from failed addition of second heart field cells to the lengthening outflow tract. Since removal of *Nedd4* in second heart field derivatives (*Mef2cAHF-Cre; Nedd4^fl/fl^*) did not lead to such outflow tract defects (Fig. 1), this suggests a novel role for *Nedd4* and cardiac neural crest cells in regulating second heart field development.

**Figure 2.**
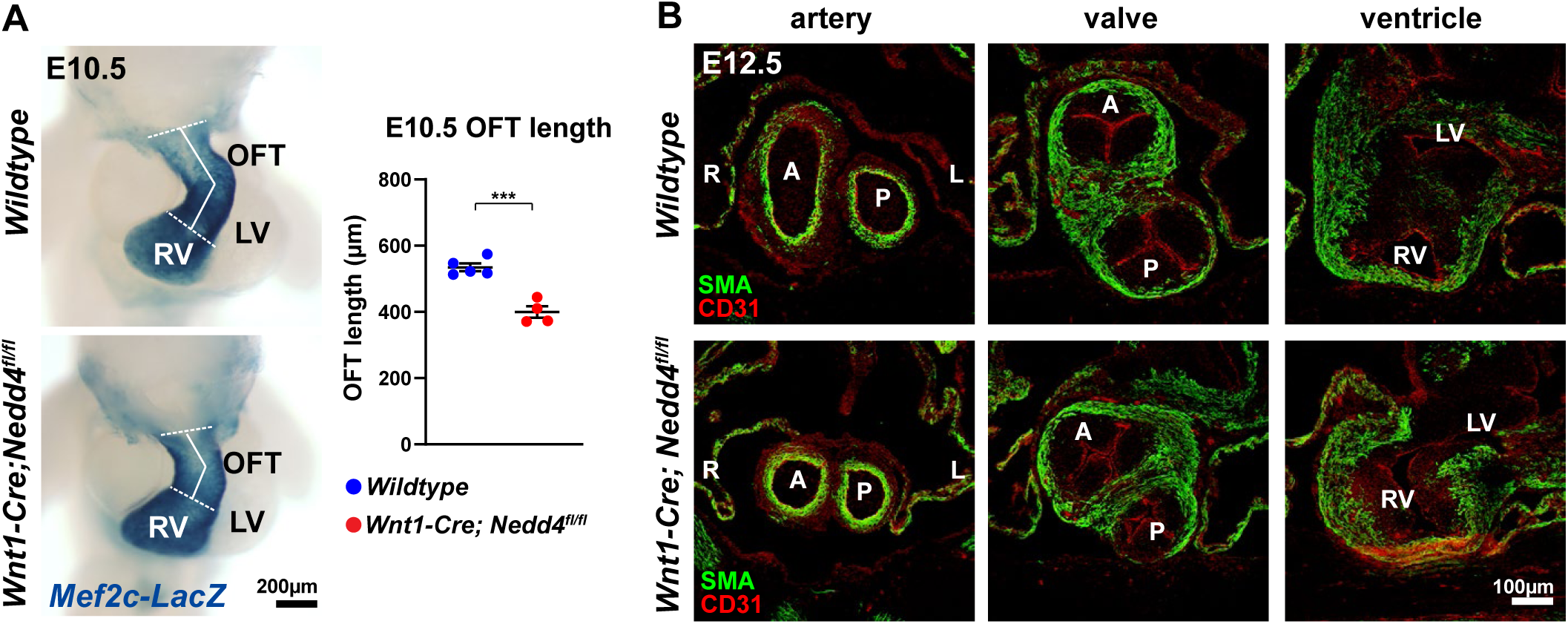
Hearts of *Wnt1-Cre; Nedd4^fl/fl^* embryos exhibit shortened outflow tract and incomplete outflow tract rotation. **A:** Hearts of *wildtype* and *Wnt1-Cre; Nedd4^fl/fl^* E10.5 embryos showing expression of *Mef2cAHF-LacZ* reporter gene in the right ventricle (RV) and outflow tract (OFT). Right panel: OFT length was measured as the distance along the dashed lines as indicated. LV, left ventricle. *** p<0.001. **B:** Coronal sections through the outflow tract region of *wildtype* and *Wnt1-Cre; Nedd4^fl/fl^* E12.5 embryos immunostained for smooth muscle actin (SMA) and CD31. Incomplete clock-wise outflow tract rotation is observed in *Wnt1-Cre; Nedd4^fl/fl^*embryos, evidenced by misalignment of the aortic valve over the right ventricle. Valve leaflet defects are also observed. L, left; R, right; A, aortic artery/valve; P, pulmonary artery/valve.

Lineage tracing of neural crest cells using an EGFP reporter revealed that prior to entering the outflow tract, cardiac neural crest cells migrate in close proximity to the second heart field in E9.5 embryos (Fig. 3A,B). This spatial relationship presents an appropriate environment for neural crest cells to interact with the second heart field during development. It is noted that *Wnt1-Cre; Nedd4^fl/fl^* embryos did not exhibit any deficiency in cardiac neural crest cells (Fig. 3A,B), nor were changes in cell proliferation or rate of cell death observed in the tissue domain populated by the neural crest cells and second heart field tissue (Supp. Fig. 1). Furthermore, expression of the second heart field marker Isl1 was unaffected (Fig. 3C and Supp. Fig. 2), as were additional markers of the second heart field and other cardiac components (Supp. Fig. 2), suggesting the development of other cardiogenic tissues was not affected by loss of *Nedd4* function in neural crest cells. Taken together, this further supports a role for disruption of *Nedd4* function in neural crest cells specifically impacting the morphogenesis of the outflow tract, and not due to loss or misspecification of either the neural crest or second heart field tissues.

**Figure 3.**
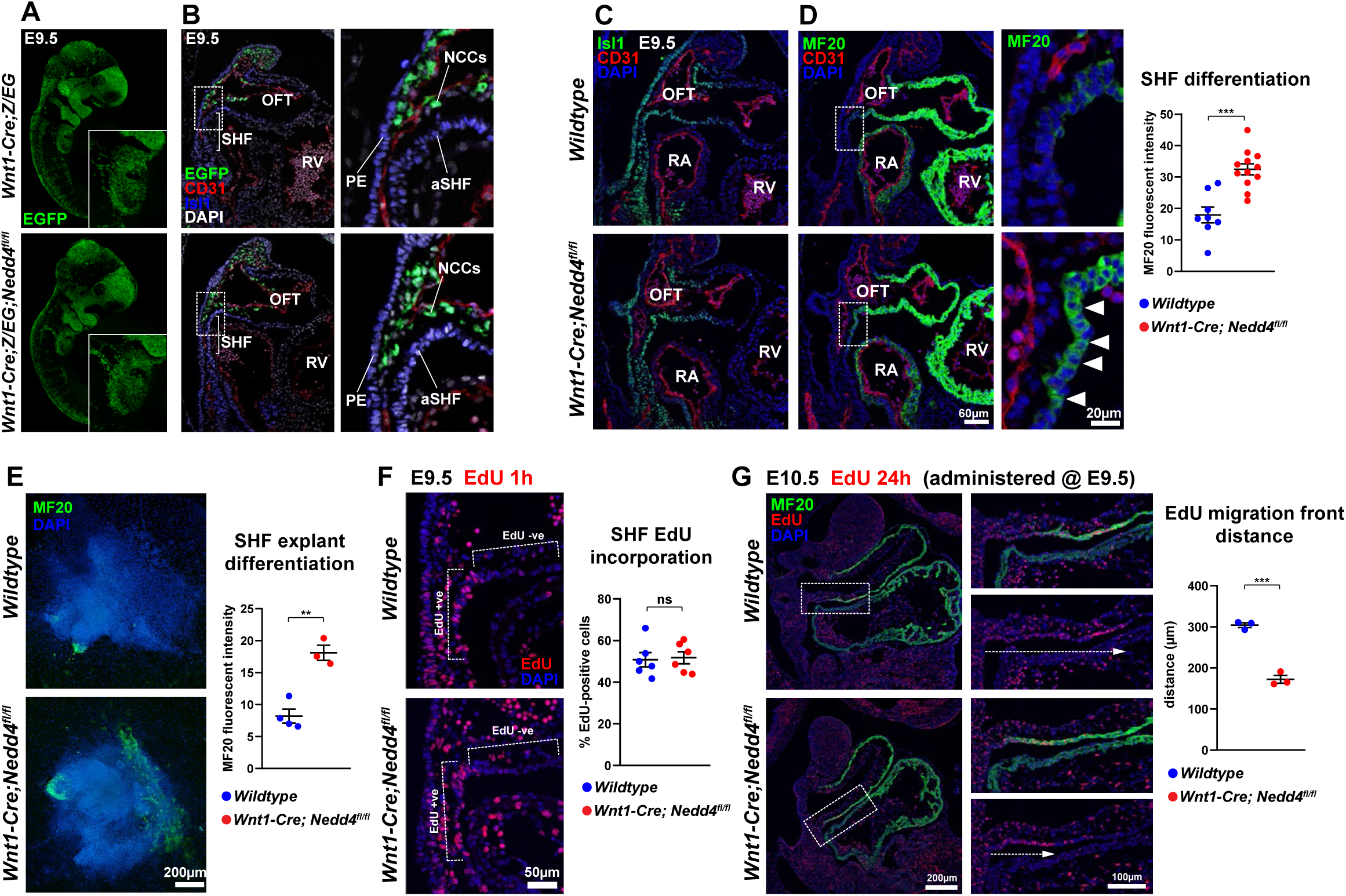
*Wnt1-Cre; Nedd4^fl/fl^* embryos display precocious differentiation and defective deployment of anterior second heart field derivatives. **A:** Whole E9.5 *Wnt1-Cre; Z/EG* and *Wnt1-Cre; Z/EG; Nedd4^fl/fl^* embryos immunostained for EGFP which labels neural crest cells and derivatives. Inset image shows the migratory cardiac neural crest cells in the 3^rd^, 4^th^ and 6^th^ pharyngeal arches. **B:** Sagittal sections through the outflow tract of E9.5 *Wnt1-Cre; Z/EG* and *Wnt1-Cre; Z/EG; Nedd4^fl/fl^*embryos immunostained for EGFP, CD31 and Isl1, revealing localisation of neural crest cells (NCCs) in close proximity to the anterior second heart field (SHF), and outflow tract (OFT). RV, right ventricle; PE, pharyngeal endoderm. **C:** Sagittal sections of E9.5 *wildtype* and *Wnt1-Cre; Nedd4^fl/fl^* embryos immunostained for Isl1 and CD31. **D:** Adjacent serial section to C immunostained for MF20 and CD31. Inset shows magnified view of anterior SHF, with arrowheads indicating precocious SHF differentiation in the mutant embryo. Right panel: Quantification of MF20 fluorescent intensity. *** p<0.001. **E:** Second heart field explant of *wildtype* and *Wnt1-Cre; Nedd4^fl/fl^* embryos, immunostained for MF20, with quantified fluorescent intensity. ** p<0.01. **F:** Sagittal sections through the outflow tract of E9.5 *wildtype* and *Wnt1-Cre; Nedd4^fl/fl^*embryos treated with EdU 1hr prior to collection. Fluorescent staining for EdU incorporation highlights low levels of EdU incorporation in the OFT, but high levels of incorporation in the highly proliferative second heart field. Labelling of EdU +ve cells in the SHF is equivalent between *wildtype* and *Wnt1-Cre; Nedd4^fl/fl^* embryos. ns, not significant. **G:** Sagittal sections through the outflow tract of E10.5 *wildtype* and *Wnt1-Cre; Nedd4^fl/fl^* embryos treated with EdU 24h prior at E9.5. EdU marks cells which have deployed from the SHF into the OFT. The migration distance of EdU+ cells was measured along the dashed arrow, and the distance quantified (right panel). *** p<0.001.

Myocardial differentiation, reported by MF20 expression, is initiated as second heart field cardiac progenitors enter the developing outflow tract. While in wildtype embryos the second heart field maintains minimal expression of MF20, *Wnt1-Cre; Nedd4^fl/fl^* embryos exhibited precocious MF20 expression in the second heart field (Fig. 3D). The enhanced MF20 expression was also observed in *ex vivo* second heart field explant cultures (Fig. 3E), supporting the inference that loss of *Nedd4* in neural crest cells causes premature differentiation of second heart field cardiac progenitors. Precocious myocardial differentiation is reputed to be one mechanism underpinning defective outflow tract lengthening, due to reducing the pool of cardiac progenitors available for contribution to the growing outflow tract ^2^. To assess effects of premature differentiation on second heart field deployment into the outflow tract, we used EdU to label the highly-proliferative second heart field (relative to the lowly-proliferative outflow tract myocardium) at E9.5, and tracked the extent of deployment of EdU +ve cells into the outflow tract 24 hours after labelling ^13^. Initial EdU labelling of the second heart field was equivalent between genotypes (Fig. 3F), while after 24h, EdU-labelled cells had moved a shorter distance into the outflow tract of *Wnt1-Cre; Nedd4^fl/fl^* embryos (Fig. 3G), consistent with impeded deployment of second heart field cells into the outflow tract underpinning outflow tract lengthening defects.

Neural crest cells have been implicated in modulating BMP and FGF signalling in the second heart field to impact outflow tract development ^14,15^. Expression of *Bmp4* and *Fgf8*, as well as downstream signalling components phospho-SMAD1/5/9 and phospho-ERK1/2, were not affected in *Wnt1-Cre; Nedd4^fl/fl^* embryos (Supp. Fig. 2 and Supp. Fig. 3). Hence, we hypothesised neural crest cells may be regulating the differentiation dynamics of the second heart field via other molecular mechanisms or signalling pathways.

### Neural crest derived DKK1 modulates canonical Wnt signalling activity in the second heart field

To investigate how loss of *Nedd4* in neural crest cells causes premature second heart field differentiation, we used laser capture microdissection to isolate cells from a defined region of the anterior second field, as well as neural crest cells and pharyngeal endoderm in the immediate vicinity, from wildtype and *Wnt1-Cre; Nedd4^fl/fl^* embryos for transcriptome profiling (Fig. 4A and Supp. Dataset 1). 897 differentially expressed genes (DEGs) with p<0.05 were identified (Fig. 4B). Gene ontology enrichment analysis of the top 30 DEGs (Fig. 4C) revealed Wnt signalling as the top signalling pathway with differential gene expression (Supp. Fig. 4A,B). *Dkk1*, the gene encoding a secreted Wnt signalling antagonist, was the top DEG identified (Fig. 4C). We next performed immunostaining and *in situ* hybridisation to investigate cell-type specificity of gene expression. Immunostaining revealed DKK1 expression was localised exclusively to neural crest cells (that co-express the neural crest marker AP2α) in the peri-second heart field region (Fig. 4D). *In situ* hybridisation, in parallel with immunostaining, validated the enhanced *Dkk1* expression and enriched DKK1 abundance in cardiac neural crest cells in *Wnt1-Cre; Nedd4^fl/fl^* embryos (Fig. 4E-G). Given the function of DKK1 as a canonical Wnt signalling inhibitor, and the proximity of cardiac neural crest cells to the anterior second heart field, we hypothesised that neural crest-derived DKK1 may antagonise Wnt signalling in the adjacent second heart field. Employing β-catenin protein stabilisation and nuclear translocation as a readout of canonical Wnt signalling activity ^16^, immunostaining for β-catenin revealed a specific reduction in nuclear localised staining intensity in the anterior second heart field and outflow tract myocardium in *Wnt1-Cre; Nedd4^fl/fl^*embryos, while staining in the pharyngeal endoderm and right atrium appeared unaffected (Fig. 4H). As well as signalling functions, β-catenin is also involved in regulating cell-adhesion via interactions with cadherins and the actin cytoskeleton ^17^. Recent work has demonstrated requisite roles for cell-adhesion, polarity and epithelial-like properties of the second heart field in outflow tract development ^18,19^. Immunostaining for markers of cell-adhesion, polarity and ECM components did not reveal any changes of these cellular attributes in the second heart field region of *Wnt1-Cre; Nedd4^fl/fl^* embryos (Supp. Fig. 5). Taken together, our data supports a role for neural crest-derived DKK1 in modulating canonical Wnt signalling activity in the second heart field.

**Figure 4.**
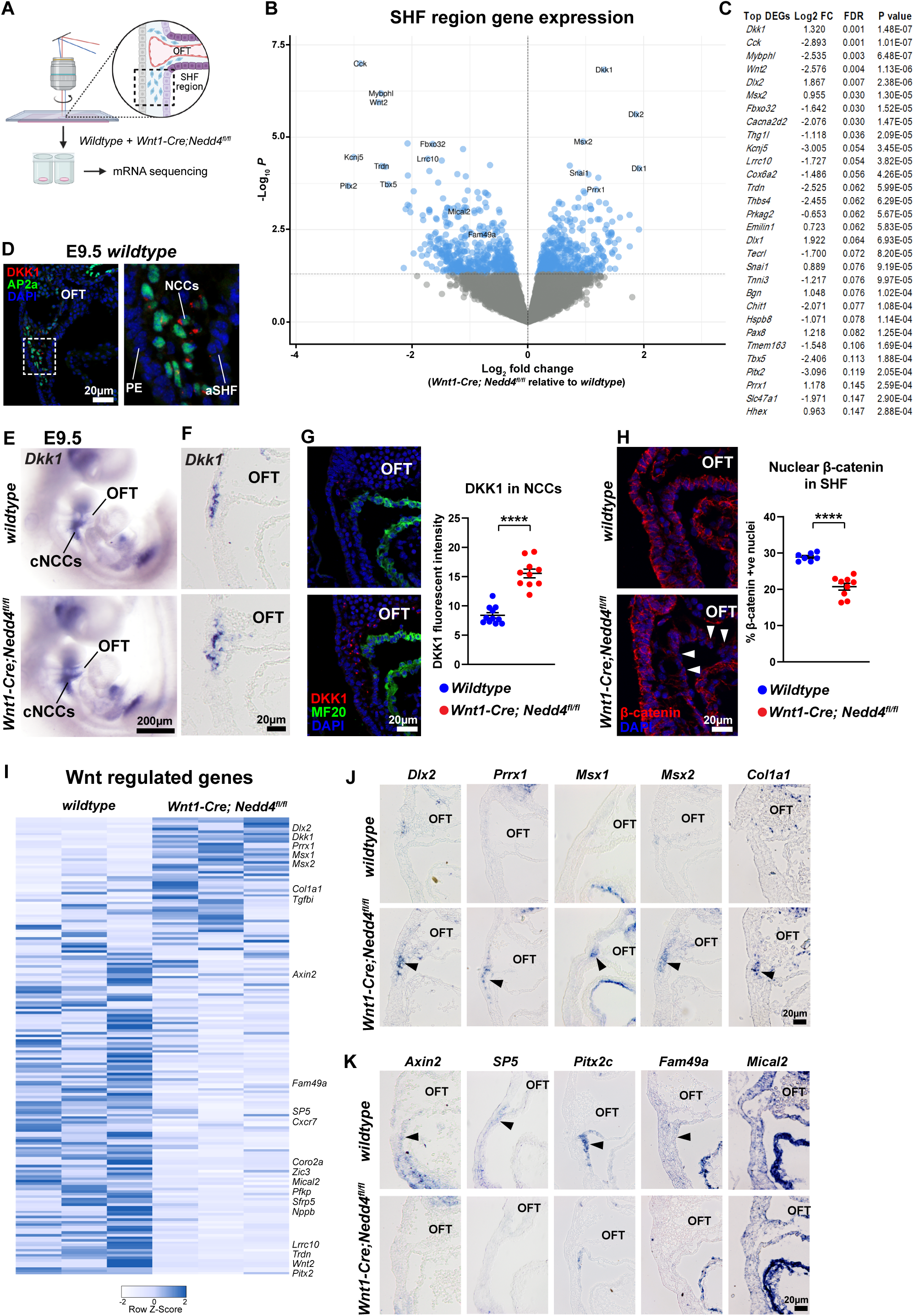
Laser capture mRNA sequencing reveals disrupted Wnt signalling in *Wnt1-Cre; Nedd4^fl/fl^* embryos. **A:** Diagram indicating tissue region (dashed box) that was excised by laser capture for mRNA sequencing, from n=3 *wildtype* and *Wnt1-Cre; Nedd4^fl/fl^* embryos. **B:** Volcano plot representing differentially expressed genes (DEGs) in *Wnt1-Cre; Nedd4^fl/fl^* tissue samples relative to *wildtype.* **C:** List of top 30 DEGs identified from laser capture mRNA sequencing. **D:** Sagittal section through the outflow tract of an E9.5 *wildtype* embryo immunostained for DKK1 and AP2α which marks the nucleus of cardiac neural crest cells (cNCCs). Inset: higher magnification demonstrates DKK1 is expressed in neural crest cells, but not in anterior second heart field (aSHF) or pharyngeal endoderm (PE). **E:** Wholemount *in situ* hybridisation of E9.5 *wildtype* and *Wnt1-Cre; Nedd4^fl/fl^* embryos, showing *Dkk1* expression in cardiac NCCs. **F:** *In situ* hybridisation of E9.5 *wildtype* and *Wnt1-Cre; Nedd4^fl/fl^* embryo sagittal sections showing *Dkk1* expression in cardiac NCCs. OFT, outflow tract. **G:** Sagittal sections of E9.5 *wildtype* and *Wnt1-Cre; Nedd4^fl/fl^* embryos immunostained for DKK1 and MF20, highlighting marked increased expression of DKK1 in NCCs in *Wnt1-Cre; Nedd4^fl/fl^* embryos, and quantified fluorescence intensity. **** p<0.0001. **H:** Sagittal sections of E9.5 *wildtype* and *Wnt1-Cre; Nedd4^fl/fl^* embryos immunostained for β-catenin. Arrowheads mark reduced β-catenin in the SHF and myocardium of the OFT. Nuclear β-catenin staining in the SHF is quantified. **** p<0.0001. **I:** Heat map of differentially expressed Wnt regulated genes. **J:** *In situ* hybridisation of E9.5 *wildtype* and *Wnt1-Cre; Nedd4^fl/fl^*embryo sagittal sections, showing increased expression of Wnt regulated genes in *Wnt1-Cre; Nedd4^fl/fl^* embryos. Arrowhead indicates expression in neural crest cells. **K:** *In situ* hybridisation of *Wnt1-Cre; Nedd4^fl/fl^* embryos showing diminished expression of Wnt regulated genes. Arrowhead indicates expression in second heart field tissue.

To assess canonical Wnt signalling activity in *Wnt1-Cre; Nedd4^fl/fl^*embryos, we intersected our laser-capture tissue mRNAseq dataset with a β-catenin ChIPseq dataset ^20^ to analyse expression of canonical Wnt target genes. We identified 179 differentially expressed Wnt target genes, with 49 upregulated and 130 downregulated (Fig. 4I). To validate and determine cell-type specificity of expression, we performed *in situ* hybridisation for ten candidate genes on tissue sections of the outflow tract region from E9.5 *wildtype* and *Wnt1-Cre; Nedd4^fl/fl^* embryos. Downregulated Wnt target genes, eg. *Axin2, SP5, Pitx2c*, *Fam49a, Mical2*, were localised to the second heart field (Fig. 4K), consistent with the reduction in nuclear β-catenin in the second heart field. Of note, upregulated expression of Wnt target genes was observed in the cardiac neural crest cells (Fig. 4J), suggesting there may be a cell-autonomous role for *Nedd4* in regulating Wnt activity in neural crest cells.

### Disruption of canonical Wnt signalling activity underpins outflow defects

To ascertain that disruption of Wnt signalling activity causes outflow tract defects in *Wnt1-Cre; Nedd4^fl/fl^* embryos, we manipulated canonical Wnt signalling *in vivo* by *in utero* treatment with small molecule activators (CHIR99021) and inhibitors (iCRT3) of Wnt signalling over the developmental time-period when neural crest cells interact with the second heart field (Fig. 5A). Treatment with Wnt agonist CHIR99021, which may enhance Wnt signalling activity, did not rescue the outflow tract defects in *Wnt1-Cre; Nedd4^fl/fl^* embryos by E13.5 (Fig. 5B,C), suggesting that escalating Wnt signalling activity could not counteract the effect of elevated DKK1 activity. However, when Wnt signalling was blocked with the chemical inhibitor iCRT3, *Wnt1-Cre; Nedd4^fl/+^* heterozygous embryos, which do not normally exhibit structural heart defects, now exhibited outflow tract rotation defects (Fig. 5B,C). Thus, loss of one copy of *Nedd4* in neural crest cells sensitises the phenotypic response to disruption of Wnt signalling, and points to genetic interaction between *Nedd4* and the Wnt signalling pathway.

**Figure 5.**
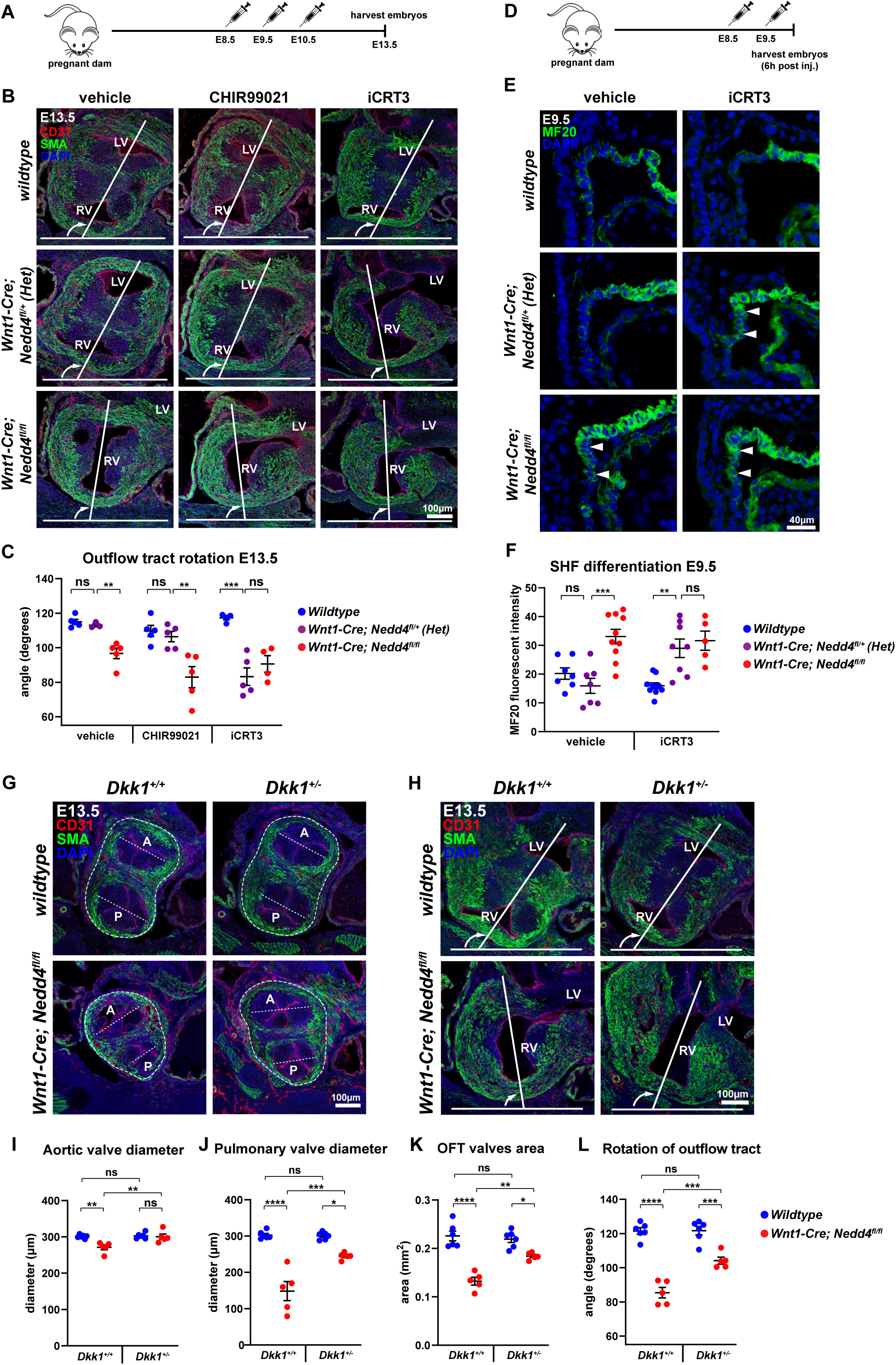
Reduced canonical Wnt signalling underpins precocious second heart field differentiation. **A:** Protocol for drug administration at E8.5, E9.5 and E10.5, and sample collection at E13.5. **B:** Coronal sections through the outflow tract region of E13.5 *wildtype*, *Wnt1-Cre; Nedd4^fl/+^* and *Wnt1-Cre; Nedd4^fl/fl^* embryos immunostained for smooth muscle actin (SMA) and CD31. RV and LV indicate positions of right ventricle and left ventricle, respectively. Outflow tract rotation was measured as the degree of clockwise rotation, indicated from the curved arrow from a perpendicular reference plane (the transverse plane of the embryo). **C:** Quantitation of outflow tract rotation. *Wnt1-Cre; Nedd4^fl/fl^* embryos exhibit deficient rotation in all treatment conditions, while *Wnt1-Cre; Nedd4^fl/+^* embryos also exhibit deficient rotation only when treated with the canonical Wnt signalling inhibitor iCRT3, and are indistinguishable from *Wnt1-Cre; Nedd4^fl/fl^*embryos. **D:** Protocol for drug administration at E8.5 and E9.5 and sample collection 6 hours after last treatment. **E:** Sagittal sections through the outflow tract region of E9.5 *wildtype*, *Wnt1-Cre; Nedd4^fl/+^*and *Wnt1-Cre; Nedd4^fl/fl^* embryos immunostained for MF20. Arrowheads indicate precocious SHF differentiation. **F:** Quantitation of MF20 fluorescent intensity in the SHF. *Wnt1-Cre; Nedd4^fl/+^* embryos exhibit precocious SHF differentiation when treated with iCRT3, and are indistinguishable from *Wnt1-Cre; Nedd4^fl/fl^* embryos. **G:** Coronal sections through the outflow tract valves of E13.5 *wildtype* and *Wnt1-Cre; Nedd4^fl/fl^* embryos crossed to *Dkk1^+/-^*and immunostained for SMA and CD31. A, aortic valve; P, pulmonary valve. Dashed lines represent valvular diameter and OFT tissue area measured. **H:** Coronal sections through the outflow tract region of E13.5 *wildtype* and *Wnt1-Cre; Nedd4^fl/fl^* embryos crossed to *Dkk1^+/-^*and immunostained for SMA and CD31. Outflow tract rotation was measured as in (B). **I-K:** Quantitation of aortic and pulmonary valve diameter, and outflow tract area as indicated in (G). These measurements are either fully or partially restored in *Wnt1-Cre; Nedd4^fl/fl^*; *Dkk1^+/-^* embryos. **L:** Quantitation of outflow tract rotation. Rotation is partially restored in partially restored in *Wnt1-Cre; Nedd4^fl/fl^*; *Dkk1^+/-^* embryos. * p<0.05; ** p<0.01; *** p<0.001; **** p<0.0001; ns, not significant.

To determine if the outflow tract rotation deficiency in iCRT3-treated *Wnt1-Cre; Nedd4^fl/+^* heterozygous embryos is caused by premature second heart field differentiation, we examined the differentiation of second heart field tissue in E9.5+6h iCRT3-treated embryos by MF20 immunostaining (Fig. 5D). iCRT3-treated *Wnt1-Cre; Nedd4^fl/+^* heterozygous embryos exhibited precocious second heart field differentiation, like that in *Wnt1-Cre; Nedd4^fl/fl^* knockout embryos (Fig. 5E,F). This finding suggests that a reduced level of canonical Wnt signalling underpins premature second heart field differentiation, and is causative of outflow tract rotation deficiency.

To determine if ectopic *Dkk1* overexpression is a contributing factor of outflow tract defects, we generated compound *Wnt1-Cre; Nedd4^fl/fl^*; *Dkk1^+/-^* embryos in which *Dkk1* expression was reduced in the *Wnt1-Cre; Nedd4^fl/fl^* background, to test for genetic rescue of outflow tract defects. Compared to *Wnt1-Cre; Nedd4^fl/fl^*embryos, *Wnt1-Cre; Nedd4^fl/fl^*; *Dkk1^+/-^* embryos exhibited improvement in outflow tract development at E13.5 (Fig. 5G,H), including restored of aortic valve diameter (Fig. 5I) partially rescued pulmonary valve diameter (Fig. 5J) and partial rescue of outflow tract tissue mass (Fig. 5K). While artery-ventricle alignment was not completely rescued in *Wnt1-Cre; Nedd4^fl/fl^*; *Dkk1^+/-^* embryos, there was a greater extent of outflow tract rotation (Fig. 5L). Taken together, the enhanced outflow tract development and rotation suggest that ectopic *Dkk1* overexpression is one major factor underpinning defective outflow tract development in *Wnt1-Cre; Nedd4^fl/fl^* embryos.

### DKK1 is a substrate for NEDD4-mediated ubiquitination

Expression of DKK1, at both transcript and protein levels, was elevated in neural crest cells with loss of *Nedd4* function (Fig. 4). Given NEDD4 is an E3 ubiquitin ligase with important roles in regulating protein abundance, we investigated if DKK1 may be a ubiquitinated substrate of NEDD4. Computational analysis on the UbiBrowser bioinformatic platform ^21^ predicts DKK1 can be ubiquitinated, with NEDD4 predicted to be the targeting E3 ligase. Using *in vitro* ubiquitination assays, we demonstrated that wildtype NEDD4 indeed ubiquitinates DKK1, while an inactive version of NEDD4 in which the catalytic cysteine is mutated (NEDD4 CS) is unable to ubiquitinate DKK1 (Fig. 6A and Supp. Fig. 6A). Moreover, NEDD4 specifically ubiquitinates DKK1, but not other DKK family members such as DKK2 (Fig. 6B). This ubiquitination likely targets DKK1 for protein degradation, as overexpression of NEDD4 WT in HEK293T cells reduces the steady-state levels of DKK1-Myc, whereas overexpression of NEDD4 CS did not affect DKK1-Myc levels (Fig. 6C). Overexpression of DKK1-GFP and FLAG-NEDD4 in HeLa cells revealed an inverse correlation of fluorescence intensity that measured protein abundance between DKK1 and NEDD4 protein, but correlation was observed between DKK1 and NEDD4 CS (Fig. 6D and Supp. Fig. 7A). Expression of a GFP-only construct with NEDD4 WT and NEDD4 CS revealed no inverse correlation (Fig. 6E), suggesting that DKK1 is the substrate targeted by NEDD4. Hence, the increased levels of DKK1 in the neural crest cells of *Wnt1-Cre; Nedd4^fl/fl^* embryos can be accounted for in part by the absence of NEDD4-mediated ubiquitination and consequent degradation of DKK1.

**Figure 6.**
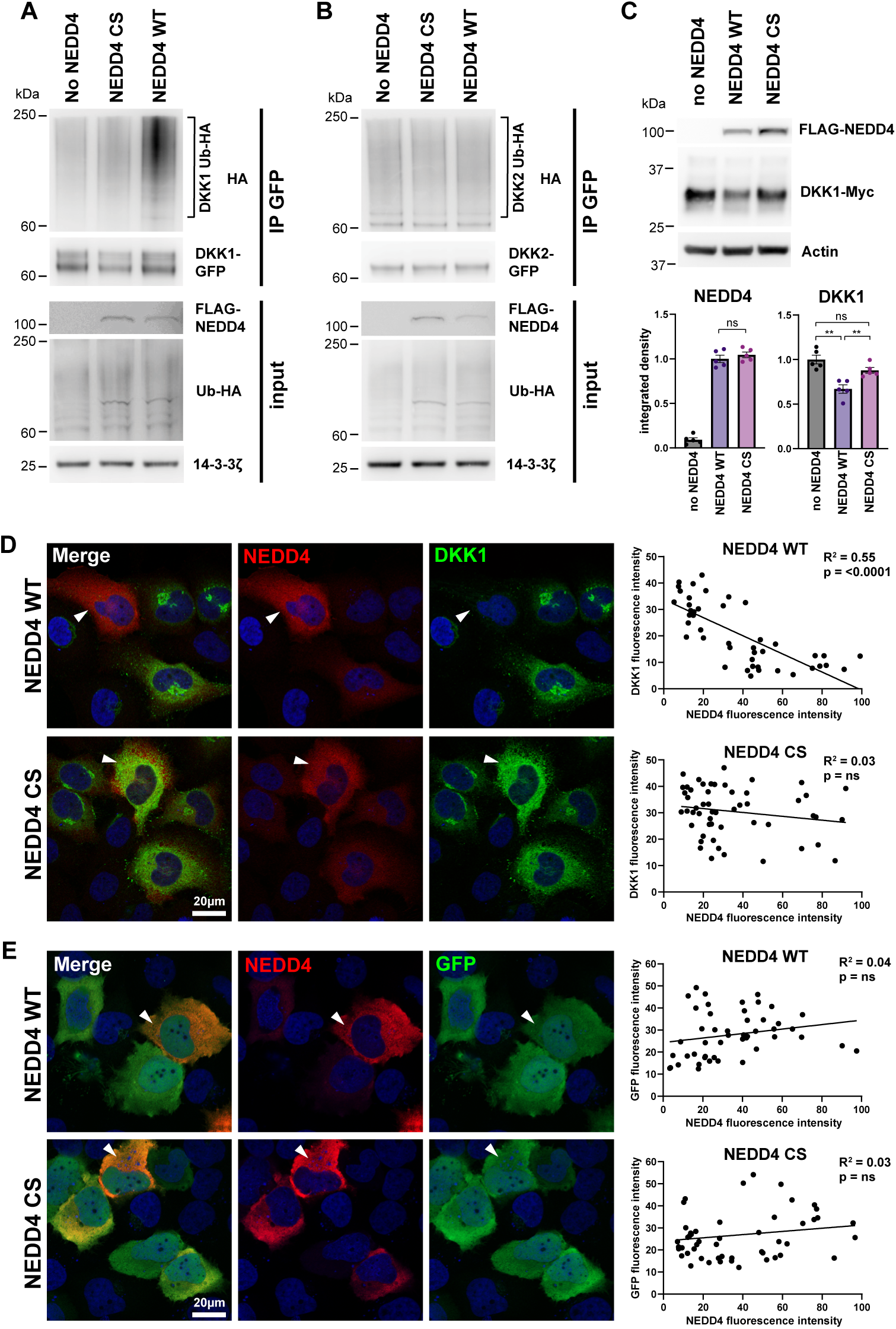
NEDD4 ubiquitinates DKK1 to regulate protein levels. **A:** Western blot of in vitro ubiquitination assay. HEK293T cells were transfected with DKK1-GFP, Ub-HA and either no NEDD4, FLAG-NEDD4 CS (cysteine mutant) or FLAG-NEDD4 WT (wildtype). Immunoprecipitation for GFP demonstrates HA positivity with NEDD4 WT, indicating ubiquitination of DKK1, but minimal ubiquitination with NEDD4 CS or no NEDD4. **B:** Ubiquitination assay as in A, but with DKK2-GFP construct. Minimal ubiquitination is observed in all conditions. **C:** Western blot of HEK293T cells transfected with DKK1-Myc and FLAG-NEDD4 as indicated. Quantitation of integrated density normalised to Actin shows that steady-state DKK1 protein levels are reduced with NEDD4 WT compared to either no NEDD4 or NEDD4 CS. ns, not significant; ** p<0.01. **D:** HeLa cells transfected with DKK1-GFP and FLAG-NEDD4 WT or CS, immunostained for FLAG and DKK1. Mean fluorescence intensity of DKK1 and NEDD4 quantified for individual cells and plotted graphically shows a negative linear correlation between DKK1 and NEDD4 WT levels, but not NEDD4 CS. For transfection control for DKK1 construct, see Supp. Fig. 7. **E:** HeLa cells transfected with GFP and FLAG-NEDD4 WT or CS, immunostained for FLAG and GFP. Quantification of mean fluorescence intensity reveals no correlation between NEDD4 and GFP levels.

### NEDD4 loss-of-function variant is clinically implicated in congenital heart defects

We identified a homozygous missense variant in *NEDD4* in a child with Tetralogy of Fallot. The child and parents were recruited and assessed as part of an Australian congenital heart disease genome sequencing study ^22^ (now extended to n = 363 trios and singletons). Analysis of genome sequencing data from the trio revealed the recessive inheritance of a *NEDD4* variant by the proband from parents carrying one allele each (Fig. 7A). The variant NEDD4:c.2297A>G:p.K766R (GenBank:NM_006154.4) changes the reference lysine to an arginine, which is predicted to be damaging by *in silico* metrics CADD and PolyPhen (CADD: 25, PolyPhen 0.995). No predicted damaging variants were identified in other genes presently known to cause congenital heart disease, making the *NEDD4* variant a compelling candidate for causing disease in this family. The parents were not consanguineous. While the family also reported congenital heart disease in distant relatives on the maternal side of the family, no clinical data were available.

**Figure 7.**
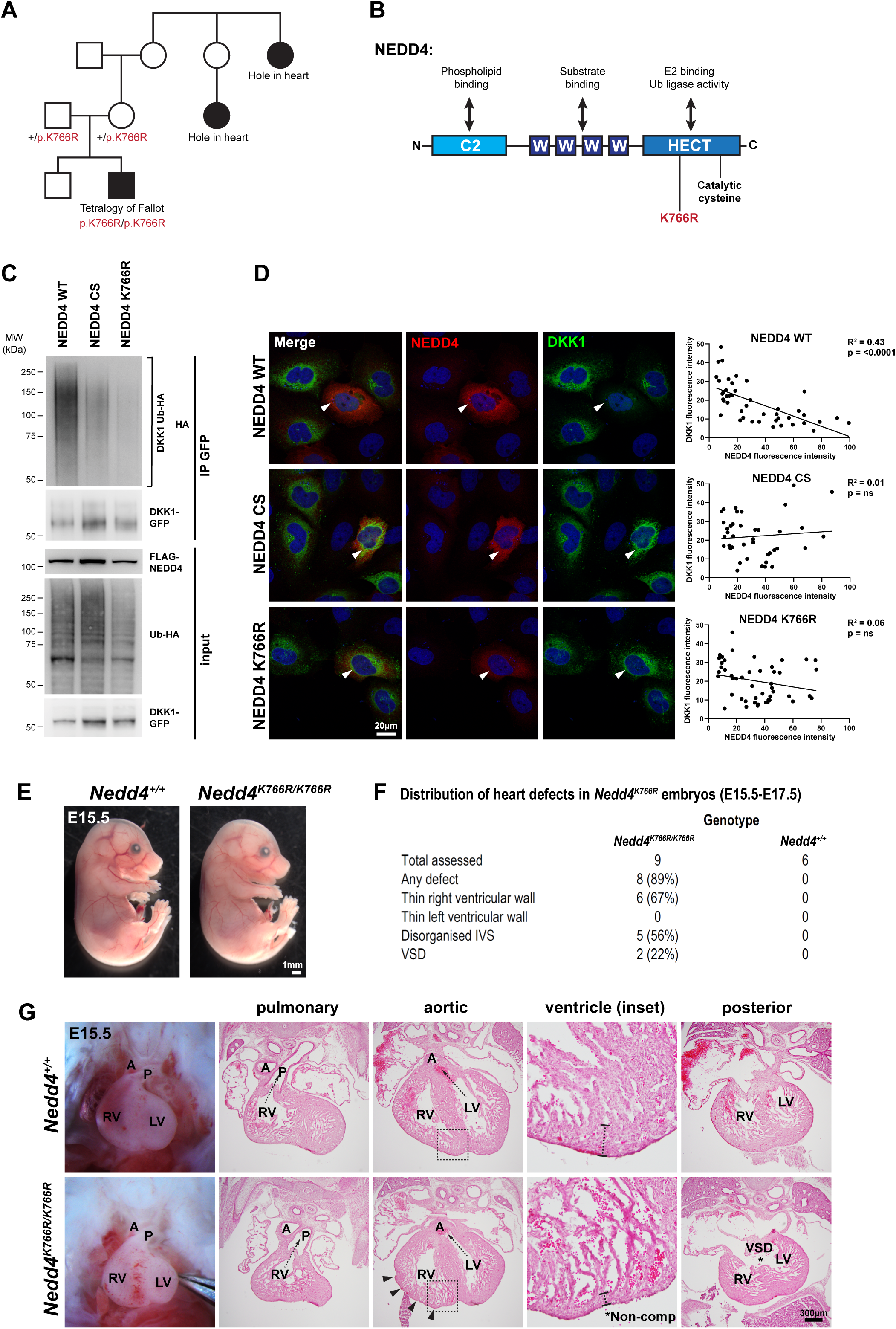
A *NEDD4* variant identified in an individual with congenital heart disease has impaired ubiquitination DKK1. **A:** Pedigree of family with a *NEDD4* variant. Genomic sequencing of the trio (parents and proband) revealed a homozygous *NEDD4* missense variant c.2297A>G (p.K766R) in the proband with Tetralogy of Fallot. Hole in the heart was the term used by the family to describe distant relatives, with no clinical data available. **B:** Protein domain structure of NEDD4, indicating the position of the K766R amino acid substitution. **C:** Western blot of in vitro ubiquitination assay. HEK293T cells were co-transfected with DKK1-GFP, Ub-HA and either NEDD4 WT, FLAG-NEDD4 CS (cysteine mutant) or FLAG-NEDD4 K766R. Immunoprecipitation for GFP demonstrates HA positivity with NEDD4 WT, indicating ubiquitination of DKK1, but minimal ubiquitination with NEDD4 CS or NEDD4 K766R. **D:** HeLa cells transfected with DKK1-GFP and FLAG-NEDD4 WT, CS, or K766R, immunostained for FLAG and DKK1. Mean fluorescence intensity of DKK1 and NEDD4 were quantified for individual cells and plotted graphically. Simple linear regression analysis reveals a negative correlation between DKK1 and NEDD4 WT levels, but not for NEDD4 CS or K766R. For transfection control for DKK1 construct, see Supp. Fig. 7. ns, not significant. **E:** Representative image of control and *Nedd4^K766R/K766R^*E15.5 embryos. **F:** Frequency of heart defects in E15.5-E17.5 *Nedd4^K766R/K766R^* embryos, with defects noted in right ventricle and interventricular septum. **G:** Histological sections of control and *Nedd4^K766R/K766R^*E15.5 embryos showing rugged surface of right ventricular wall (arrowheads), right ventricular non-compaction, and ventricular septal defect (VSD).

The K766R amino acid change is located in the HECT domain of NEDD4, which is responsible for ubiquitin ligase activity (Fig. 7B). Given that NEDD4 can ubiquitinate DKK1, *in vitro* assays to assess the effect of K766R amino acid substitution on NEDD4 function revealed NEDD4 K766R had reduced capacity to ubiquitinate DKK1 (Fig. 7C and Supp. Fig. 6B). Furthermore, analysis of fluorescence intensity revealed no correlation between DKK1 and NEDD4 K766R protein levels (Fig. 7D and Supp. Fig. 7B), confirming NEDD4 K766R is a loss-of-function variant for ubiquitination of DKK1.

To investigate the *in vivo* consequences of NEDD4 K766R substitution, we generated a CRISPR-engineered *Nedd4^K^*^766^*^R/K766R^* mouse model. Homozygous embryos of this mutant mouse strain were present at expected mendelian ratios at all developmental stages and phenotypically normal (Fig. 7E). Examination of hearts in *Nedd4^K766R/K766R^* homozygous embryos revealed a range of developmental cardiac defects (Fig. 7F). While artery-ventricle alignment defects such as overriding aorta, which is a clinical feature of Tetralogy of Fallot, were not present, there were defects consistent with deficiency of right ventricular development. This included a thin ventricular wall which was restricted to the right ventricle, often with a rugged appearance (Fig. 7G) suggestive of defective right ventricular compaction, a disorganised interventricular septum and ventricular septal defects (Fig. 7G). Hence *Nedd4^K766R/K766R^* embryos exhibit heart defects including some features of Tetralogy of Fallot such as right ventricular wall and interventricular septal defects, implicating the NEDD4 K766R variant as a cause of congenital heart disease.

Taken together, NEDD4 regulation of DKK1 in neural crest cells unveils a novel mechanistic paradigm in heart development, which when disrupted leads to heart defects in mice and humans.

## Discussion

Wnt signalling plays critical roles in multiple aspects of heart development, where it has stage-dependent positive and negative effects on progenitor cell specification, maintenance and differentiation (reviewed in ^23,24^). Canonical Wnt/β-catenin signalling is required for initial mesoderm induction in the gastrulating embryo, however it is then downregulated to promote cardiac precursor specification which concomitantly is supported by activation of non-canonical Wnt signalling ^25,26^. As heart development progresses, cells of the second heart field require canonical Wnt/β-catenin signalling to maintain cells in a proliferative and undifferentiated state. As second heart field progenitors move into the outflow tract, canonical Wnt signalling is gradually downregulated, arresting cell proliferation and initiating myocardial differentiation, in which non-canonical Wnt signalling plays an important role. Hence, the temporal balance of activation and inhibition of Wnt signalling is essential for orchestrating the proliferation and differentiation dynamics of cardiac progenitors during heart development.

Conditional knockout and constitutive activation of β-catenin in specific cardiac cell lineages support a role for the dynamic regulation of canonical Wnt signalling in heart development. Knockout of β-catenin in the *Islet1-Cre* or *SM22α-Cre* lineages results in failure of the right ventricle to form, while overactivation of β-catenin causes right ventricular enlargement ^27,28^. In other lineages (*Mef2cAHF-Cre* and *MesP1-Cre*), both loss- and gain-of-function of β-catenin are detrimental to right ventricle formation ^29,30^. Hence, the precise dosage and timing of Wnt/β-catenin signalling is critical. Inactivation of β-catenin target genes in the second heart field also support a role for canonical Wnt signalling in outflow tract development. For example, conditional knockout of *Pitx2* in *Mef2cAHF-Cre* and *Islet1-Cre* lineages causes artery-ventricle alignment defects including double outlet right ventricle and transposition of the great arteries ^31^, and is consistent with the phenotype of *Wnt1-Cre; Nedd4^fl/fl^* embryos which exhibit downregulation of *Pitx2c* in the second heart field.

While Wnt signalling is known for its role in second heart field development, the source of Wnt ligands has remained unclear. A recent study suggests that Wnt2 secreted from the first heart field can promote second heart field proliferation at early stages of heart development (E8.5) ^32^. Whether Wnt2 is the key Wnt ligand responsible for maintaining the second heart field in a proliferative state at later stages (E9.5) during the critical time period of outflow tract lengthening remains to be determined. Nevertheless, as anterior second heart field cells move into the outflow tract, canonical Wnt signalling is downregulated to promote myocardial differentiation. This reduction in Wnt signalling may be a consequence of these cells moving away from a localised source of Wnt morphogen as they enter the outflow tract. Alternatively, we propose cardiac neural crest cells can act as a critical rheostat of Wnt signalling as second heart field cells transition into the outflow tract. Neural crest cells are spatially positioned in this transition zone; providing a source of DKK1 to fine tune Wnt signals and promote coordinated second heart field differentiation.

Neural crest cells have previously been implicated in second heart field development. Early studies in avian models demonstrated that surgical ablation of the premigratory cardiac neural crest caused an array of conotruncal heart defects ^4^, underpinned by failed addition of second heart field cells to the lengthening outflow tract ^6,7^. Furthermore, these studies showed loss of cardiac neural crest cells led to increased FGF signalling and excessive proliferation in the second heart field, suggesting cardiac neural crest act to modulate FGF signalling to promote appropriate differentiation of the outflow tract myocardium ^5,33,34^. BMP signalling antagonises the pro-proliferative effects of FGF signalling to promote myocardial differentiation, and neural crest cells have been shown to be essential for mediating the effects of BMP on cardiomyocyte differentiation ^14,15^. Disruption of downstream BMP signalling components in neural crest cells also causes abnormal second heart field differentiation and outflow tract defects ^35,36^. How neural crest cells mediate BMP-induced second heart field differentiation is unknown, however an attractive hypothesis is that BMP signalling may induce neural crest cells to secrete another factor which in turn supresses FGF signalling and/or promotes differentiation. Wnt/β-catenin signalling also feeds into these complex signalling interactions, with Wnt gain- and loss-of-function also affecting BMP and FGF signalling outcomes in the second heart field ^28–30^. As no disruption of FGF and BMP signalling, or changes in proliferation of second heart field progenitors were observed in *Wnt1-Cre; Nedd4^fl/fl^* embryos, this may suggest neural crest-derived factors act downstream of these signalling interactions in this model to regulate myocardial differentiation of the second heart field. Hence, FGF and BMP signals may be essential to prime the second heart field to appropriately respond to differentiation signals, with neural crest cells acting as a critical source of DKK1 to modulate Wnt signalling and actively initiate myocardial differentiation.

DKK1 is ideally located to act as a rheostat of Wnt signalling, providing precise spatial and temporal control of canonical Wnt signalling activity in the second heart field. This dynamic regulation is essential for maintaining a critical balance of second heart field progenitor maintenance versus differentiation, to foster appropriate outflow tract elongation. Our study has unveiled expression of DKK1 specifically in cardiac neural crest cells, providing new mechanistic data to explain how neural crest cells regulate second heart field myocardial differentiation. Enhanced DKK1 activity induces cardiac differentiation in embryonic stem cell models ^27,37^, and in Xenopus embryos ^38^, which is consistent with the increased myocardial differentiation observed in the second heart field of *Wnt1-Cre; Nedd4^fl/fl^* embryos. While *Dkk1* knockout mouse embryos have no reported heart defects ^39^, *Dkk1* and *Dkk2* double knockout embryos exhibit mild heart defects including ventricular septal defects and myocardial/epicardial hyperplasia ^40^. Findings of our present study are supportive of ectopic gain-of-function of DKK1, rather than loss-of-function, as being causative of developmental defects of the heart. The impact of widespread ectopic gain-of-function of *Dkk1* on heart development remains to be investigated.

Our study reports the ubiquitin modification of the DKK1 protein as an additional level of post-translational control to modify DKK1 protein abundance, and hence further regulates the precise level of Wnt signalling activity. Our finding provides a molecular mechanism linking loss of the ubiquitin ligase NEDD4 in neural crest cells to changes in DKK1 abundance. While we have focussed on post-translational regulation of DKK1 protein, we also observe an increase in *Dkk1* mRNA in neural crest cells of *Wnt1-Cre; Nedd4^fl/fl^* embryos. Indeed, many of the differentially expressed genes that were upregulated in *Wnt1-Cre; Nedd4^fl/fl^* embryos are localised to the cardiac neural crest. This likely points to cell-autonomous roles for *Nedd4* loss of function in cardiac neural crest cells, which may also underpin cardiac defects. This would be a subject of future studies in heart development.

The clinical significance of our findings is highlighted by the outcome of the functional study of a deleterious homozygous missense variant of *NEDD4* in an individual with Tetralogy of Fallot. This NEDD4 variant results in a non-synonymous amino acid change in the protein domain responsible for ubiquitin ligase activity, and we show that this variant has greatly reduced ability to ubiquitinate DKK1, as well as exhibiting heart defects in a CRISPR engineered *in vivo* model. The clinical Tetralogy of Fallot phenotype overlaps with the heart defect phenotype of *Wnt1-Cre; Nedd4^fl/fl^* mice, implicating an association of neural crest-derived NEDD4 and congenital heart disease. Mis-regulation of DKK1 activity may represent a new pathogenic mechanism underpinning congenital heart disease. Neural crest cells have never-before been considered as paracrine modifiers of Wnt signalling activity. We here define a new developmental paradigm in heart morphogenesis, and implicate neural crest cell modulation of Wnt signalling as a key mechanistic attribute controlling heart development.

## Methods

### Mice

All experiments were carried out in accordance with ethical guidelines of the University of South Australia Animal Ethics Committee. *Nedd4^-/-^* mice and *Nedd4^fl/fl^* have been described previously ^41,42^. To remove *Nedd4* in specific tissues, we crossed *Nedd4^fl/fl^* females to *Wnt1-Cre;Nedd4^fl/+^*^43^, *Mef2cAHF-Cre;Nedd4^fl/+^*^44^ or *Tie2-Cre;Nedd4^fl/+^* ^45^ males. *Wnt1-Cre; Nedd4^fl/fl^* mice were intercrossed with *Dkk1^+/-^*^39^ mice to generate *Wnt1-Cre; Nedd4^fl/fl^*; *Dkk1^+/-^* embryos for assessing the rescue effect of DKK1 dosage compensation. The following reporter lines were also intercrossed with *Wnt1-Cre;Nedd4^fl/+^* mice; *Mef2cAHF-LacZ* ^46^ and *Z/EG* ^47^. *Nedd4^K766R^* mice were generated using CRISPR homology directed repair. SaCas9 KKH was used with a single stranded DNA donor template oligo with 60bp homology arms either side of the K766R site and a silent CTC(L764) > TTG(L764) mutation to abolish the PAM site and introduce a BclI recognition site for genotyping. To obtain mouse embryos of defined gestational ages, mice were mated in the evening and the morning of vaginal plug formation was counted as E0.5.

### Drug treatments of pregnant dams

Pregnant dams were intraperitoneally injected with 16.6 mg/kg CHIR99021 (Selleck Chem) or 10 mg/kg iCRT3 (Selleck Chem) at the indicated time points in 15% DMSO, 30% PEG 300, PBS vehicle. Embryos were harvested at E13.5 or E9.5, and processed for immunostaining.

### Histology and immunostaining

Embryos were fixed in 4% paraformaldehyde in PBS. Cryosections were stained with Hematoxylin and Eosin to classify heart defects at E15.5-E17.5. For immunolabelling, cryosections, whole embryos or fixed cells were blocked in 10% DAKO block, 0.2% BSA, 0.2% Triton X-100 in PBS, and stained with the indicated primary antibodies. Antibodies used were mouse anti-alpha smooth muscle actin (Sigma) 1:2000; rat anti-CD31 (Biolegend) 1:150; chicken anti-GFP (Millipore) 1:1000; mouse anti-Isl1 (DSHB 40.3A4) 1:50; mouse anti-MF20 (DSHB) 1:100; goat anti-DKK1 (R&D Systems) 1:200; mouse anti-AP2α (DSHB 3B5) 1:20; rabbit anti-β-catenin (Cell Signaling Technology 19807) 1:200; mouse anti-FLAG (Sigma) 1:1000; rabbit anti-phospho-Histone H3 (Millipore) 1:500; rabbit anti-cleaved-Caspase-3 (Cell Signaling Technology) 1:500; rabbit anti-phospho-SMAD1/5/9 (Cell Signaling Technology) 1:200; rabbit anti-phospho-ERK1/2 (Cell Signaling Technology) 1:100; goat anti-Scribble (Santa Cruz Biotechnology) 1:50; rabbit anti-Laminin (Sigma) 1:1000; rabbit anti-Fibronectin (DakoCytomation) 1:1000; mouse anti-N-cadherin (Cell Signaling Technology) 1:100; Alexa Fluor 647 conjugated Phalloidin (Invitrogen) 1:200. EdU staining was performed following manufacturers recommendations (Invitrogen Click-iT EdU Alexa Fluor 555 Cell Proliferation Kit). TUNEL staining was performed following the manufacturers recommendations (Roche *In Situ* Cell Death Detection Kit, TMR Red). Slides were mounted in Fluoro-mount G with DAPI (ProSciTech). Confocal images were acquired on a LSM 800 (Zeiss) system. Mean fluorescence intensity was measured using ZEN 3.4 software (Zeiss). For MF20 measurements in second heart field at E9.5, a line of 100μm was drawn from downwards perpendicular to the outflow tract, and the single layer of cells of the second heart field (pericardial wall) was traced around only to the end of the 100μm line. Hence, an equivalent area of MF20 fluorescent intensity was measured across different embryo samples.

### In situ hybridisation

Whole embryos or cryosections were hybridised with digoxigenin-labelled antisense probes for the following genes *Dkk1, Dlx2, Prrx1, Msx1, Msx2, Col1a1, Axin2, SP5, Pitx2c, Fam49a, Mical2, Tbx1, Tbx20, Isl1, HoxB1, Fgf8, Bmp4.* Signal was detected using anti-DIG-AP conjugated Fab fragments (Roche), and staining with NBT/BCIP (Roche). Whole embryos were imaged with Olympus CZX10 and tissue sections were imaged with Olympus IX73.

### β-Galactosidase Staining

Fixed embryos were incubated in staining solution: 19mM Sodium dihydrogen phosphate, 81mM Disodium hydrogen phosphate, 2mM MgCl_2_, 5mM EGTA, 0.01% Sodium deoxycholate, 0.02% NP-40, 5mM Potassium ferricyanide, 5mM Potassium ferrocyanide and 1mg/ml X-gal substrate, at 37°C until blue staining was sufficient.

### SHF explant

The second heart field region was dissected from E9.5 embryos by removing the head above pharyngeal arch 2, cutting off the outflow tract and heart, cutting down the lateral sides of the pharynx to create a tissue flap, and then separating this second heart field tissue flap at the posterior end from the rest of the embryo. This was transferred to tissue culture dishes coated with 50μg/ml Collagen and 50μg/ml Fibronectin, and cultured for 5 days in 5% FCS + DMEM. Explants were fixed in 4% PFA.

### SHF deployment assay

Pregnant dams were intraperitoneally injected with 100 mg/kg EdU (5-ethynyl-2’-deoxyuridine) at E9.5, and harvested either 1h or 24h later. Embryos or tissue sections were stained with Click-iT EdU Kit AlexaFluor-555 conjugated (Life Technologies) following staining with primary antibodies.

### Laser capture and RNA preparation

Tissue preparation and laser capture was based on ^48^. Briefly, mouse embryos were fresh-frozen in OCT, cryosectioned and collected on PEN membrane slides. Slides were stained with 1% Cresyl Violet to visualise tissue sections, airdried, and then laser dissection performed promptly (Leica LMD) to specifically capture the central anterior second heart field region (with associated neural crest and pharyngeal endoderm tissue). RNA was prepared from tissue by lysing in 4M guanidine isothiocyanate solution (Invitrogen) and precipitating with sodium acetate and ethanol. After 75% ethanol wash, RNA pellet was resuspended in 12μl H_2_O, with 1μl reserved for Bioanalyser quality control of RNA, and 9.5μl used for subsequent cDNA synthesis.

### RNA sequencing and bioinformatics analyses

Isolated RNA samples were submitted to the ACRF Genomics Facility for library preparation and sequencing. Extracted RNA was quantified using the Bioanalyser Pico chip and 9.5ul (containing 5-17ng of RNA) was input into SMART-Seq v4 Ultra Low Input RNA Kit for Sequencing (Takara Bio) for generation and amplification of double-stranded full-length cDNA (using 9 cycles of LD-PCR). Prior to library preparation, cDNA was sheared to 200-500 bp fragments using the Covaris system. NGS libraries were prepared using the ThruPLEX DNA-seq Kit (Takara Bio) together with the DNA Unique Dual Index Kit (Takara Bio), using 11 cycles of PCR for library amplification. Indexed libraries from three biological replicates for each *wildtype* and *Wnt1-Cre; Nedd4^fl/fl^*samples were pooled equally for sequencing using 2×75bp reads on an Illumina NextSeq High Output kit. Raw data, averaging 47 million reads per sample were analysed and quality checked using the FastQC program (http://www.bioinformatics.babraham.ac.uk/projects/fastqc). Reads were mapped against the mouse reference genome (mm10) using the STAR spliced alignment algorithm ^49^ (version 2.6.1d with default parameters and --chimSegmentMin 20, -- quantMode GeneCounts) returning an average unique alignment rate of 81%. The resulting mapped reads were found to possess a high rate of duplicates (≈64%). These were marked and subsequently removed using Picard Tools (version 2.16.0) ^50^ leaving, on average 18 million deduplicated reads per sample. Differential expression analysis was evaluated from TMM normalized gene counts using R (version 3.2.3) and edgeR (version 3.3) ^51^ following protocols as described ^52^. Alignments were visualised and interrogated using the Integrative Genomics Viewer v2.3.80 ^53^. Graphical representations of differentially expressed genes were generated using Glimma ^54^, Degust ^55^ and MaGIC Volcano Plot Tool ^56^. Gene ontology analysis was performed using DAVID ^57,58^.

### Ubiquitination assays

For cell-based ubiquitination assays HEK293T cells originally obtained from ATCC (CRL-3216) were grown in DMEM supplemented with 10% fetal calf serum. Plasmid transfections were carried out with Lipofectamine 2000 (Thermo Fisher). HEK293T cells were co-transfected with equal amount of FLAG-tagged NEDD4 WT, NEDD4 CS, or NEDD4 K766R, HA-tagged ubiquitin (gift from Sharad Kumar) and GFP tagged DKK1 or DKK2. The DKK1-GFP and DKK2-GFP constructs were generated by cloning a PCR fragment of the open reading frames from pCS2-DKK1-FLAG (Addgene plasmid #16690) or pCS2-DKK2-FLAG (Addgene plasmid #15495) in frame to EGFPN3 using EcoRI and BamHI restriction sites. Cells were grown for 24-48 hours at 37°C with 5% CO_2_ in a humidified atmosphere. Transfected cells were treated with 10μM MG132 and 100μM of Chloroquine for three hours prior to protein extraction.

### Immunoprecipitation and western blot

Cells were lysed in NP-40 buffer (137mM NaCl, 10mM Tris pH 7.4 (108319 Merck), 10% glycerol, 1% NP-40, 2mM sodium fluoride and 2mM sodium vanadate) containing Complete Protease Inhibitors. For purification of GFP-fusion proteins GFP-Trap beads were used following the manufacturers’ protocols (gta-10 Chromotek). Protein samples were separated by SDS-PAGE and transferred to PVDF membrane (88585 Thermo Fisher). Membranes were blocked in 5% skim milk powder in tris buffered saline (TBS, 50mM Tris pH7.5, 150mM NaCl) with 0.1% Tween 20 (P1379 Merck). The following primary antibodies were used: mouse anti-HA (6E2, Cell Signaling Technologies) 1:1000; mouse anti-FLAG (Sigma) 1:1000; mouse anti-Myc (9B11, Cell Signaling Technologies) 1:1000; rabbit anti-GFP (ClonTech) 1:1000; mouse anti-β-Actin (Sigma) 1:1000; Goat anti-DKK1 (R&D Systems) 1:500; rabbit anti 14-3-3zeta C-16 (sc1019 Santa Cruz, 1:1000). Densitometry of all western blot data was performed with ImageJ. All data are presented as mean ± SEM. All experimental studies were analysed using Student’s t test with a P value of <0.05 was considered to be statistically significant.

### HeLa protein abundance correlation assays

HeLa cells were grown in DMEM supplemented with 10% fetal calf serum. Plasmid transfections were carried out with X-tremeGENE HP (Roche) in 8-well glass bottom imaging slides (Eppendorf). Cells were co-transfected with equal amount of FLAG-tagged NEDD4 WT, NEDD4 CS, or NEDD4 K766R, and DKK1-GFP. Cells were grown for 24-48 hours at 37°C with 5% CO_2_ in a humidified atmosphere. Cells were fixed with 4% paraformaldehyde in PBS for 20 minutes, and immunostained following protocol described above. Mean fluorescent intensity for each cell was calculated using ZEN 3.4 software (Zeiss). Approximately 15 cells were quantified for each condition per experiment, for n=3 individual experiments. Graphs represent combined data for all experiments. Simple linear regression analysis was performed in GraphPad Prism.

### Human patient recruitment

Ethics approval was obtained from Sydney Children’s Hospital Network Human Research Ethics Committee (approval number HREC/16/SCHN/73). The family was recruited through the Kids Heart Research DNA Bank at Children’s Hospital at Westmead, Sydney, Australia. Informed, written consent was obtained from the family. Heart defects in the proband was confirmed by echocardiography.

### Patient sequencing and genome data analysis

Genomic DNA was extracted as described previously ^59^. DNA sample libraries were prepared using the Illumina TruSeq Nano DNA HT Library Prep Kit and genome sequenced (Illumina HiSeq X Ten, 150 base paired-end reads) at Genome.One, Garvan Institute of Medical Research, Sydney, Australia. Calling and annotation of sequencing data were performed as per ^22^. In short, sequence reads were mapped to human reference genome hg38 using Burrows-Wheeler Aligner (BWA-mem v0.7.12) ^60^. Single nucleotide variants (SNVs) and small insertions or deletions were called using Platypus v0.8.1 ^61^. The variant call files (VCFs) were annotated using ANNOVAR v 2016Feb01 ^62^. Variants were prioritised as described in ^22^ by screening for a list of curated CHD genes followed by unbiased comprehensive analysis for all genes. Variant frequencies were in reference to gnomAD database (v2.1.1) ^63^. Variant of interest identified in the family, NEDD4:c.2297A>G:p.K766R (GenBank:NM_006154.4), had an allele frequency of 0.004438.

### Statistics

In all graphical representations, individual values represent independent biological replicates. Unless otherwise specified, p values were calculated using t-test. For grouped analysis in CHIR99021 and iCRT3 treated embryos, significance was assessed using 2 way ANOVA multiple comparisons.

## Acknowledgements

This research was supported by funds from the National Health and Medical Research Council (NHMRC): Project Grant (ID1144004)[QS, SW]; Ideas Grant (ID2030340)[SW, QS]; Principal Research Fellowship (ID1135886)[SLD]; Leadership Level 3 Fellowship (ID2007896)[SLD]; Project Grant (ID1162878)[SLD, DW, EG]; Synergy Grant (ID1181325)[SLD, DW, EG]. Channel 7 Children’s Research Foundation [SW]. Royal Adelaide Hospital Mary Overton Postdoctoral Fellowship [SW]. National Heart Foundation Future Leader Fellowship [QS]. NSW Health Cardiovascular Research Capacity Program Senior Researcher Grant [SLD]. We thank Paul Riley for critical reading of the manuscript.

## Figure Legends

**Supplementary Figure 1 - *Wnt1-Cre; Nedd4^fl/fl^* embryos do not exhibit changes in cell proliferation or cell death in the second heart field region.**

**A:** Proliferation of second heart field (SHF) cells was assessed in E9.5 *wildtype* and *Wnt1-Cre; Nedd4^fl/fl^* embryos. Cryosections through the SHF region were immunostained for phospho-Histone H3 (PHH3) and Isl1. PHH3, Isl1 double-positive cells were scored and expressed as a percentage relative to total Isl1 positive cells. Counts were performed on at least 5 sections per embryo and averaged, from n=5 biological replicates. **B:** Similar sections as in A were immunostained for TUNEL and Isl1, and percentage of TUNEL, Isl1 double-positive cells calculated relative to total Isl1 positive cells. Counts were performed on at least 5 sections per embryo and averaged, from n=3 biological replicates. **C:** Similar sections as in A were immunostained for cleaved-caspase 3 and Isl1, and percentage of Casp3, Isl1 double-positive cells calculated relative to total Isl1 positive cells. Counts were performed on at least 5 sections per embryo and averaged, from n=4 biological replicates. ns, not significant.

**Supplementary Figure 2 – Heart field and cardiac development markers appear unchanged in *Wnt1-Cre; Nedd4^fl/fl^* embryos.**

Whole mount *in situ* hybridisation on E9.5 *wildtype* and *Wnt1-Cre; Nedd4^fl/fl^*embryos for transcripts as labelled.

**Supplementary Figure 3 – Phospho-SMAD and Phospho-ERK signalling appears unchanged in the second heart field region of *Wnt1-Cre; Nedd4^fl/fl^* embryos.**

**A:** Transverse sections of E9.5 *Wnt1-Cre; Z/EG* and *Wnt1-Cre; Z/EG; Nedd4^fl/fl^* embryos immunostained for phospho-SMAD1/5/9, EGFP and Isl1. **B:** Transverse sections of E9.5 *wildtype* and *Wnt1-Cre; Nedd4^fl/fl^* embryos immunostained for phospho-ERK1/2, CD31 and Isl1. NT, neural tube; PE, pharyngeal endoderm; SHF, second heart field.

**Supplementary** Figure 4 **– Gene ontology analysis of differentially expressed genes in *Wnt1-Cre; Nedd4^fl/fl^* compared to *wildtype* embryos.**

**A:** Gene ontology analysis (DAVID) of the top 30 differentially expressed genes (DEGs) from laser capture RNA-seq analysis of the second heart field region from *wildtype* and *Wnt1-Cre;Nedd4^fl/fl^*embryos. Terms assessed were “Biological process”, “Cellular component” and “Molecular function”. **B:** Gene ontology analysis of the top 100 DEGs, as in A.

**Supplementary Figure 5 – Cell polarity and extracellular matrix markers appear unchanged in the second heart field of *Wnt1-Cre; Nedd4^fl/fl^* embryos.**

**A:** Sagittal sections through the outflow tract region of E9.5 *wildtype* and *Wnt1-Cre; Nedd4^fl/fl^* embryos immunostained for Scribble and Laminin. **B:** Sagittal sections through the outflow tract region of E9.5 *wildtype* and *Wnt1-Cre; Nedd4^fl/fl^*embryos immunostained for F-actin with Phalloidin, Fibronectin and N-cadherin. OFT, outflow tract; RA, right atrium.

**Supplementary Figure 6 – Quantification of ubiquitination assays demonstrates NEDD4 ubiquitinates DKK1, but not a human disease variant NEDD4 K766R.**

**A:** Quantitation of western blots from in vitro ubiquitination assay in Figure 6A. Integrated density measurements were taken for the immunoprecipitated (IP) blots for GFP (ie. total DKK1 IP’d) and HA (ie. ubiquitinated portion of DKK1). Ubiquitinated DKK1 is expressed as the ratio of HA density measurement / GFP density measurement. Quantitation was performed for n=3 independent experiments. * p<0.05. **B:** Quantitation of western blots from in vitro ubiquitination assay in Figure 7C. Measurements performed as in A. *** p<0.001; ** p<0.01; ns, not significant.

**Supplementary Figure 7 – HeLa cell protein abundance correlation assays, additional controls.**

**A:** HeLa cells transfected with DKK1-GFP and FLAG-NEDD4 WT, NEDD4 CS, or no NEDD4 immunostained for FLAG, DKK1 and GFP, as in Figure 6D. GFP immunostaining serves as a transfection control for the DKK1-GFP construct, validating that any reduced DKK1 immunofluorescence in these cells is attributed to post-translational regulation of DKK1 by NEDD4 WT, and not simply due to lack of transfection of the DKK1-GFP construct. No NEDD4 panels serve as an additional control. **B:** Immunostaining of HeLa cells as in A, for Figure 7D, demonstrating GFP immunofluorescence for transfection control.

**Supplementary Dataset 1 – Laser capture mRNA sequencing data**

Excel file containing laser capture RNA sequencing read counts from n=3 *wildtype* (WT) and n=3 *Wnt1-Cre; Nedd4^fl/fl^* (Mut) embryos. Data includes raw counts, counts per million (CPM), differentially expressed genes (DEGs) calculated from DEGUST analysis, and Wnt regulated genes.

