## Supplementary figures for "Neural crest cell derived DKK1 modulates Wnt signalling in the second heart field to orchestrate cardiac outflow tract development"

Supplementary Figure 1

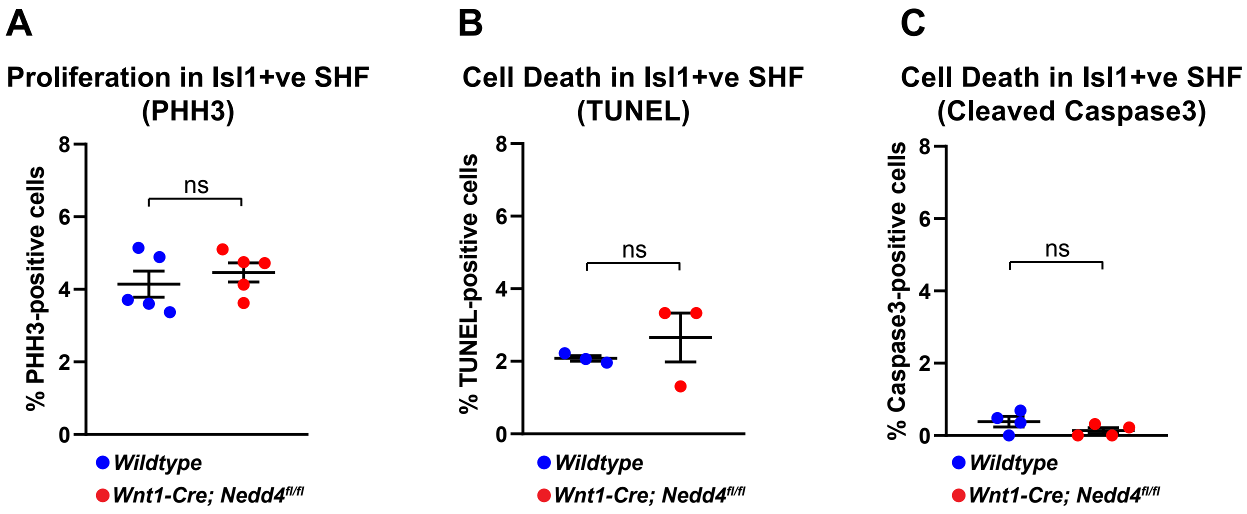

### Supplementary Figure 2

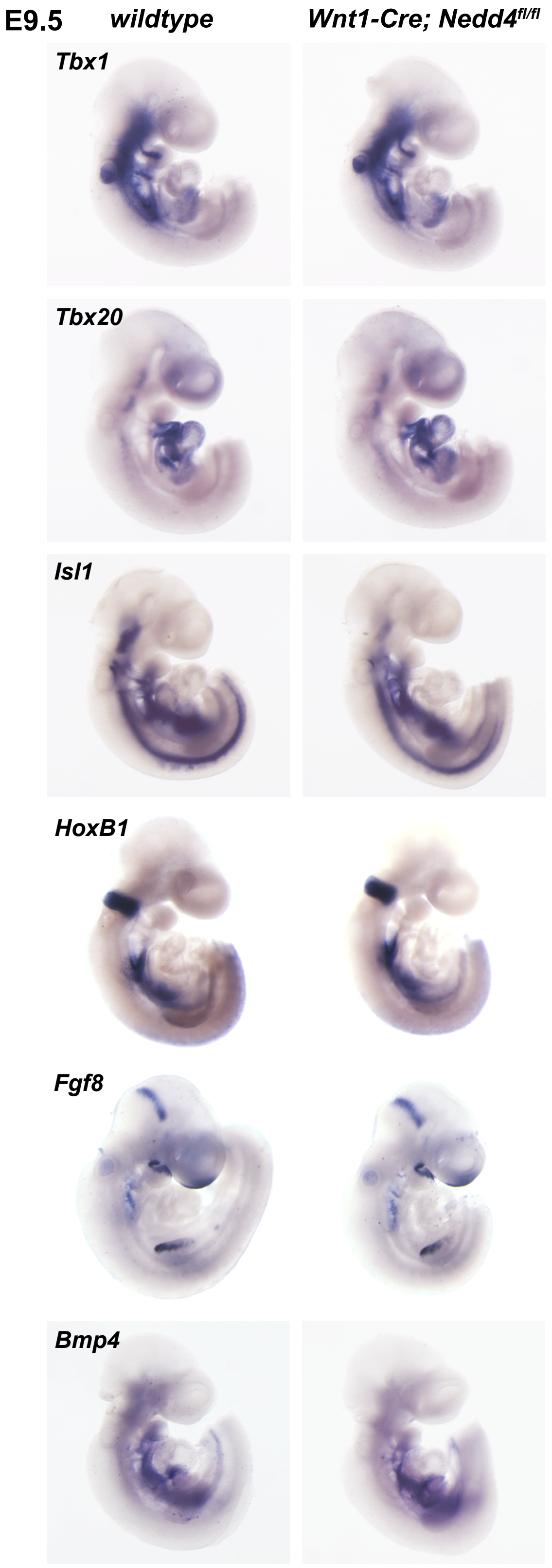

Supplementary Figure 3

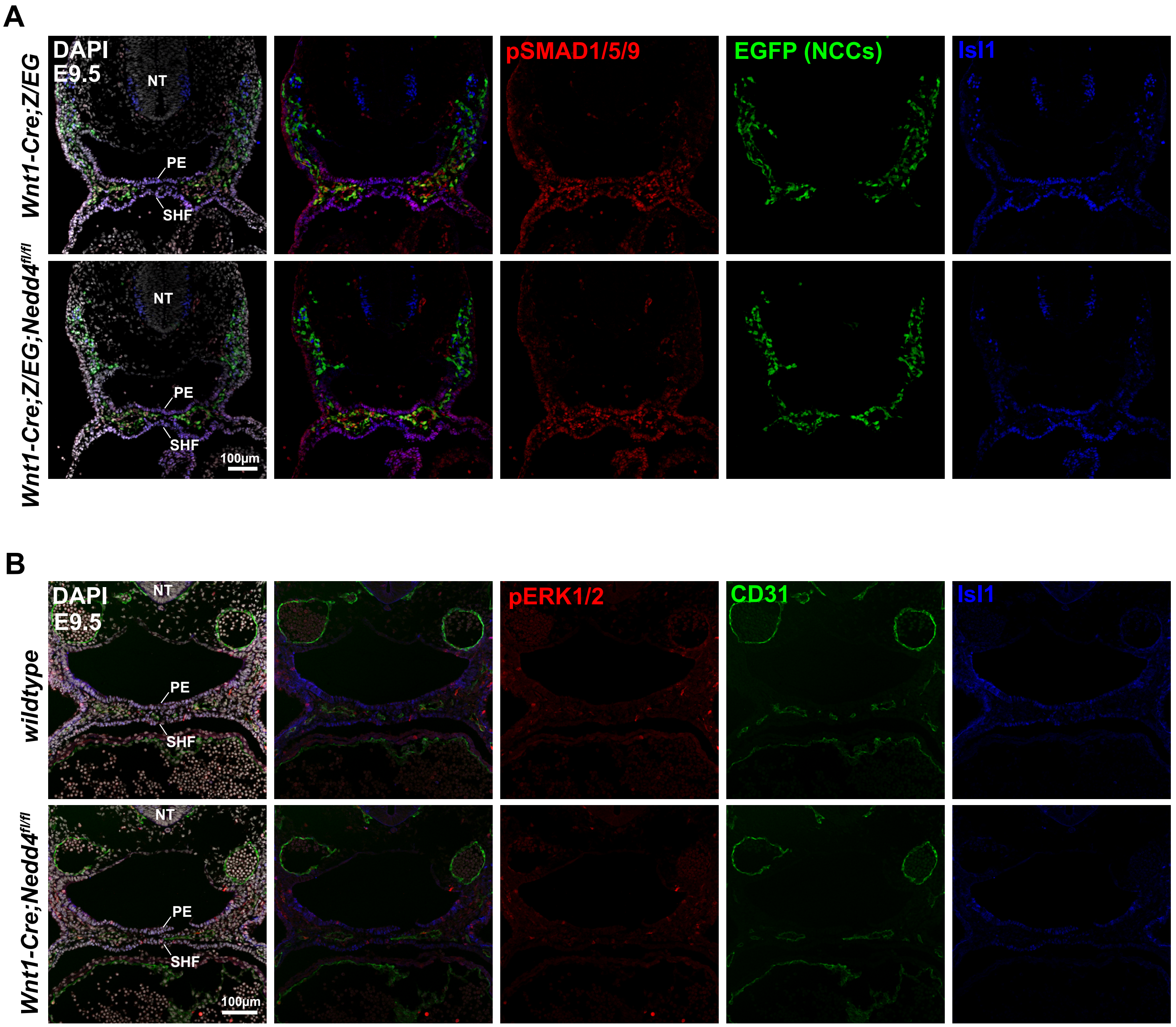

Supplementary Figure 4

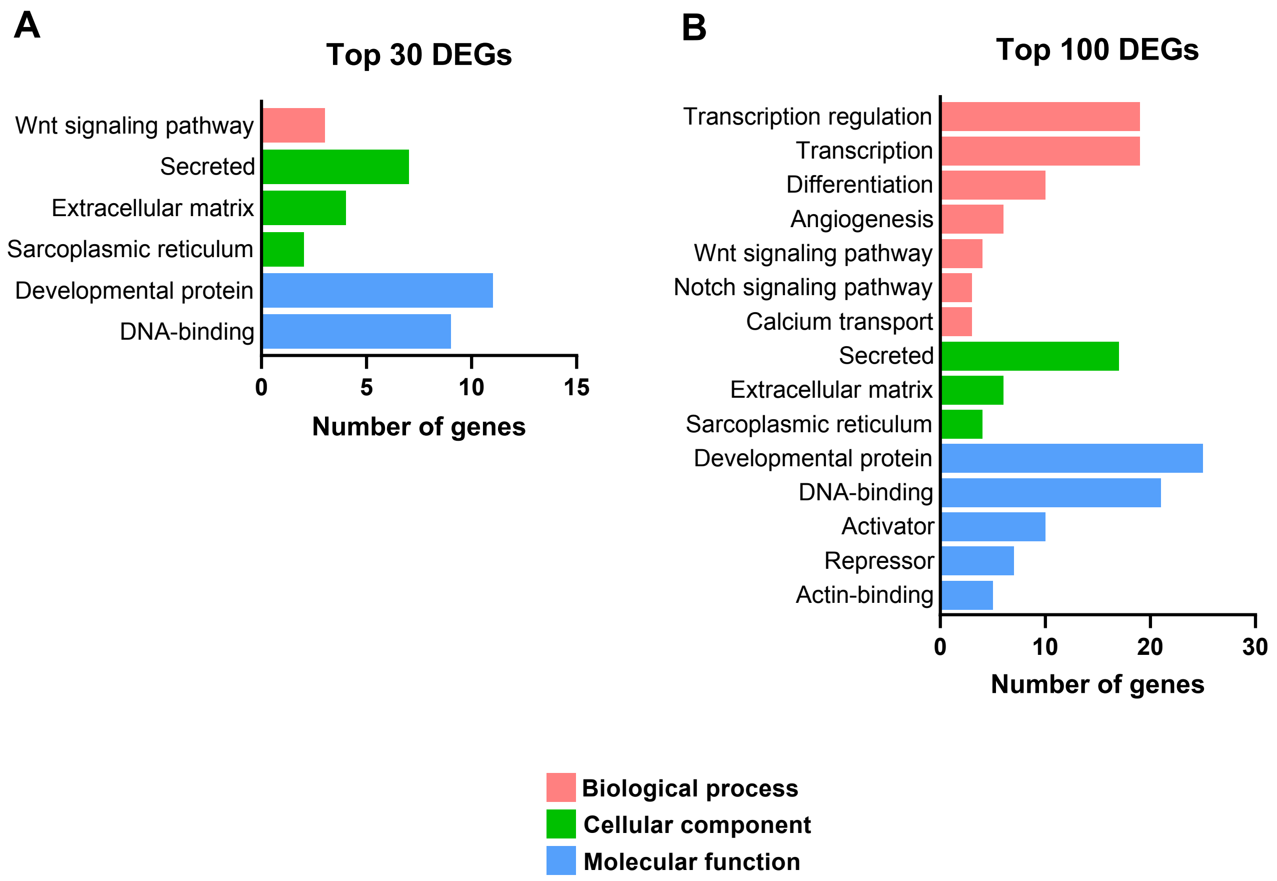

Supplementary Figure 5

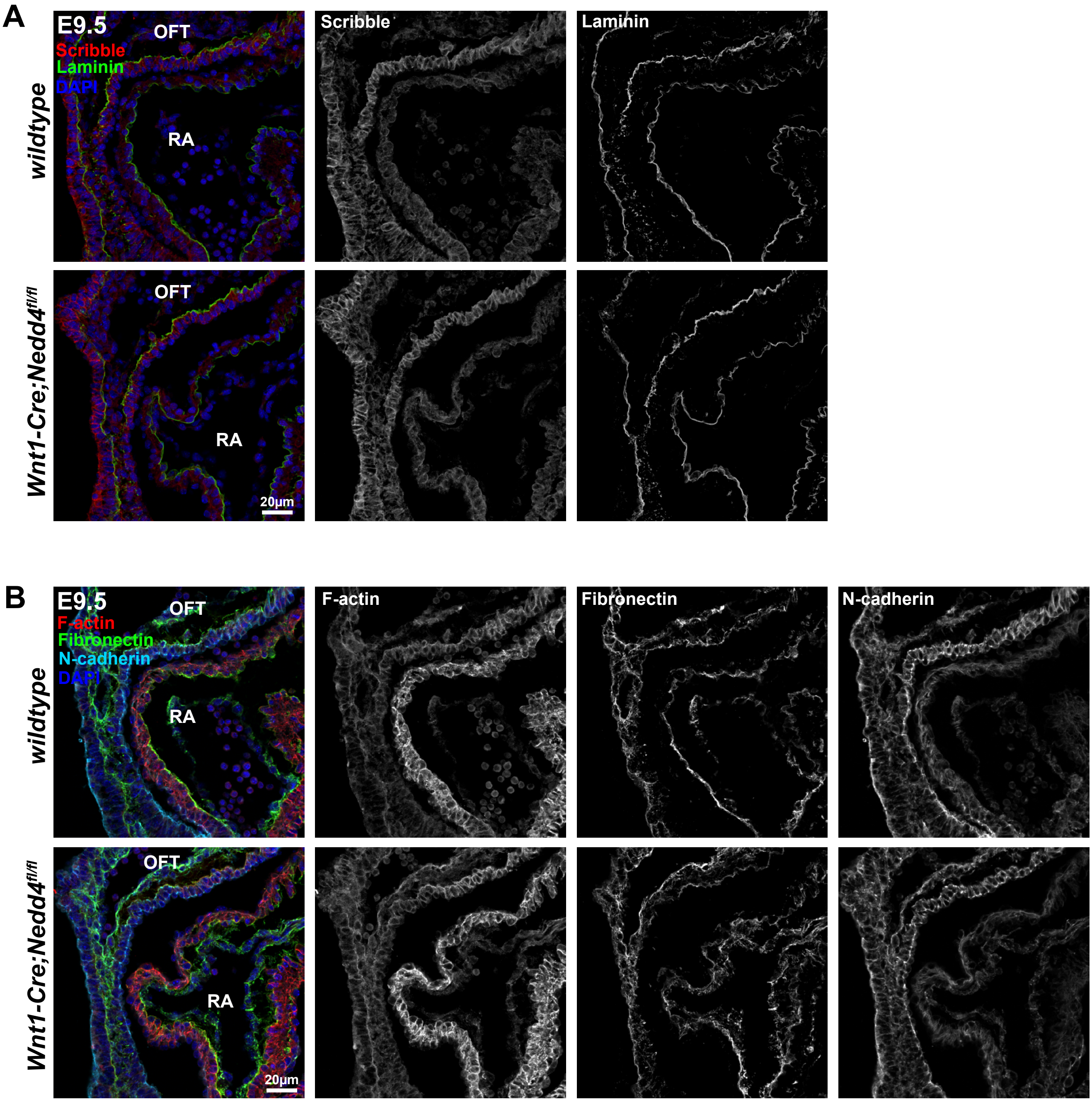

Supplementary Figure 6

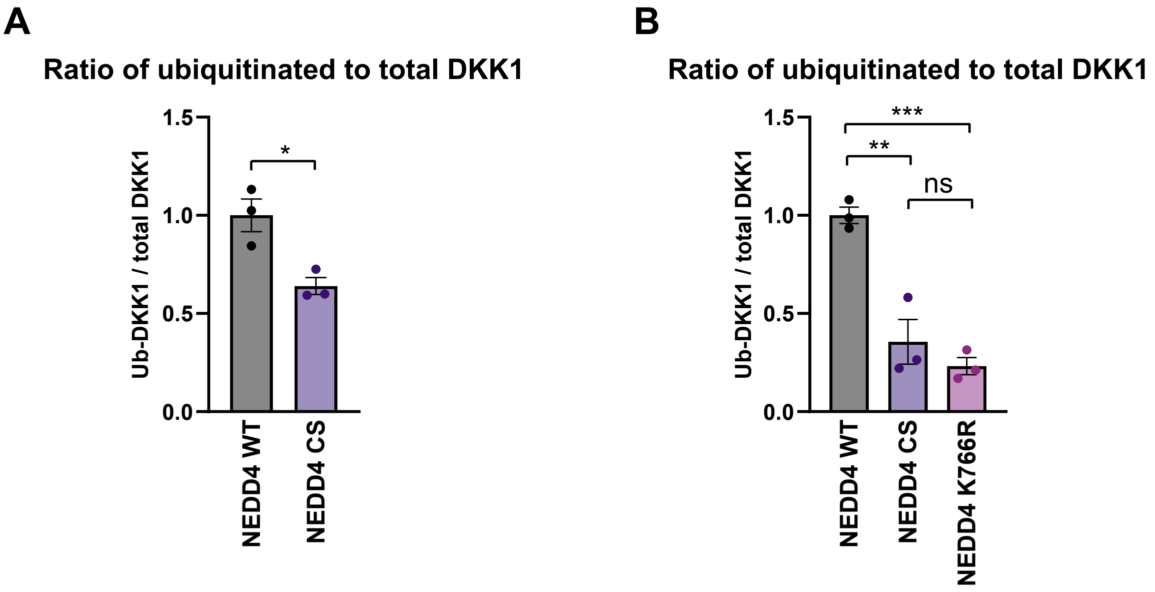

Supplementary Figure 7

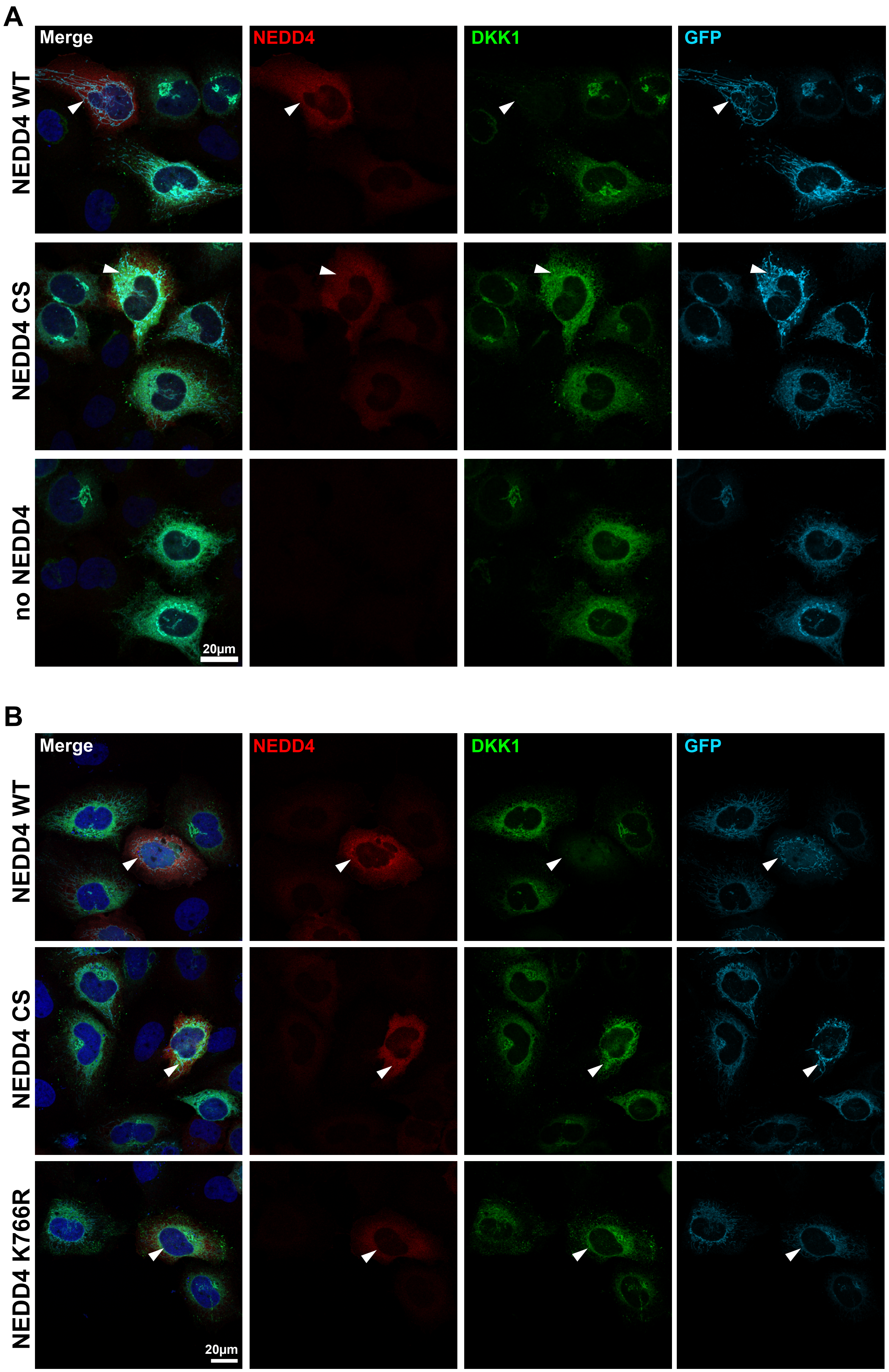
